# Chronic Prestress Regulation of Micro-Heart Muscle Physiology

**DOI:** 10.1101/2022.07.23.501265

**Authors:** Daniel W. Simmons, David R. Schuftan, Jingxuan Guo, Kasoorelope Oguntuyo, Ghiska Ramahdita, Mary K. Munsell, Brennan Kandalaft, Missy Pear, Nathaniel Huebsch

## Abstract

Engineered heart tissues have been created to study cardiac biology and disease in a setting that more closely mimics *in-vivo* heart muscle than 2D monolayer culture. Previously published studies suggest that geometrically anisotropic micro-environments are crucial for inducing “*in vivo*-like” physiology from immature cardiomyocytes. We hypothesized that such anisotropic tissue geometries regulate tissue prestress, and that in turn this prestress is a major factor regulating cardiomyocytes’ electrophysiological development. Thus, we studied the effects of tissue geometry on electrophysiology of micro-heart muscle arrays (μHM) engineered from human induced pluripotent stem cells (iPSC). Geometries predicted to increase tissue prestress not only affected cardiomyocyte structure, but also had profound effects on electrophysiology.

Elongated geometries led to adaptations that yielded increased calcium intake during each contraction cycle. Strikingly, pharmacologic studies revealed a prestress threshold is required for sodium channel function, whereas L-type calcium and rapidly-rectifying potassium channels were largely insensitive. Analysis of RNA and protein levels suggest sodium channel activity changes were related to post-transcriptional, and potentially post-translational, changes. μHM formed from Plakophilin 2 (PKP2) knockout iPSC had a cellular structure similar to isogenic controls.

However, these tissues exhibited no functional sodium current and an overall lesser degree of electrical remodeling in response to prestress. These results suggest that PKP2, a key component of the nascent desmosome, is crucial to transducing tissue prestress into physiologically beneficial electrical remodeling via activation of sodium channels.

## Introduction

Human induced pluripotent stem cell-derived cardiomyocytes (iPS-CM) are promising alternatives to current high-throughput *in-vitro* assays for predicting cardiotoxic effects of drugs, and studies on patient-derived iPS-CM have demonstrated correlations between *in-vitro* results and clinical incidence of arrhythmias.^1^ However, iPS-CM have significantly different electrophysiology:^2^ iPSC-CM have a high spontaneous beating rate, inefficient calcium handling, and altered sodium and potassium channel function, which largely persist even over long-term (up to 12 months) culture.^3, 4^ These changes cause monolayer iPS-CM to exhibit both false-positive and false-negative readouts for drug induced toxicity and pro-arrhythmia.^5–8^ These inconsistencies are likely due to deficiency in the expression and/or function of one or more ion channels.^9^

In order to overcome the limitations of monolayer iPS-CM, a variety of engineered heart tissues have been created in attempts to increase the maturity of these cells.^10–13^ Tissues provide necessary cues such as cell alignment and increased mechanical forces that have been shown to mature cells, and have been created using a wide variety of methods.^14–22^ Culturing iPS-CM within these “*in vivo*-like” formats has led to a variety of positive outcomes, including increased expression of contractile proteins, higher conduction velocity, and global changes in electrophysiology. Importantly, the resulting tissues can more accurately predict the effects of pro– and anti-arrhythmic drugs compared to monolayers, presumably through more physiologic ion channel expression and/or activity.^14–16, 20, 21, 23, 24^

While culture within engineered heart muscle formats alone is insufficient to achieve postnatal maturations states in iPS-CM, it is clear that these systems do promote iPS-CM electrical remodeling.^22^ Interestingly, different engineered tissue systems elicit differing levels of induced electrical remodeling, even absent additional biophysical (e.g. pacing^20^) or biochemical (e.g. fatty acids^25, 26^) stimuli. It remains unknown what common factor(s) these tissues contain that induce cardiomyocyte maturation, making it difficult to compare systems or determine how to further improve them.

Mechanical loading plays critical roles in regulating heart and cardiomyocyte physiology,^11^ not only during development, but also in disease progression.^27–30^ It has been shown that the amount and type of mechanical loading, including tissue geometry and prestress, regulates cardiomyocyte morphology,^31^ multiple signaling pathways, the expression of various ion channels, calcium handling, and overall animal mortality.^32^ It is thus likely that the increased levels of mechanical loading induced onto the cells in the various tissue systems leads to the changes seen in physiology.

Despite indirect evidence that mechanical forces are likely to promote electrical remodeling of iPS-CM, mechanisms through which this may occur remain unknown. Increased prestress on cardiomyocytes cultured in aligned 3D engineered tissue environments^14, 16, 20, 21, 23^ has been hypothesized as a key physical cue to induce maturation,^31^ analogous to the manner in which forces, such as prestress, regulate in-vivo heart formation and function.^28, 33, 34^ In the present study, we utilized tissues of different geometries to understand how different levels of tissue prestress regulate function of specific cardiomyocyte ion channels function to yield electrical remodeling.

## Results

### Tissue Prestress Regulates Cardiomyocyte Morphology

To test the impact of prestress in 3D tissues on electrophysiology, micro heart muscle arrays (µHM) of various geometries were created. Simple continuum mechanics models of tissue compaction, which consider isotropic shrinkage of a hyperelastic solid against anisotropic boundary conditions, predict a direct link between tissue geometry and prestress (Fig. 1A). To directly test whether these geometric changes affected the structure and function of cardiomyocytes within μHM, we used Hydrogel Assisted STereolithographic Elastomer prototyping (HASTE)^35^ to create dogbone-shaped molds from 3D-printed resins with three different geometries. These consisted of a progressively lengthened “shaft” connecting “knobs” on either end, with the extreme case being a simple knob, which our models predict would yield 3D tissues without prestress concentrations. These resulting molds were seeded with iPSC-CM at day 15 of differentiation^36^ (μHM day 15/μD15; Fig. 1B). Gross tissue morphology was largely unaffected by tissue shaft length within dogbone-shaped designs, although µHM with longer shafts exhibited slightly more lateral compaction compared to 1mm µHM (Fig. 1C, D).

**Figure 1.**
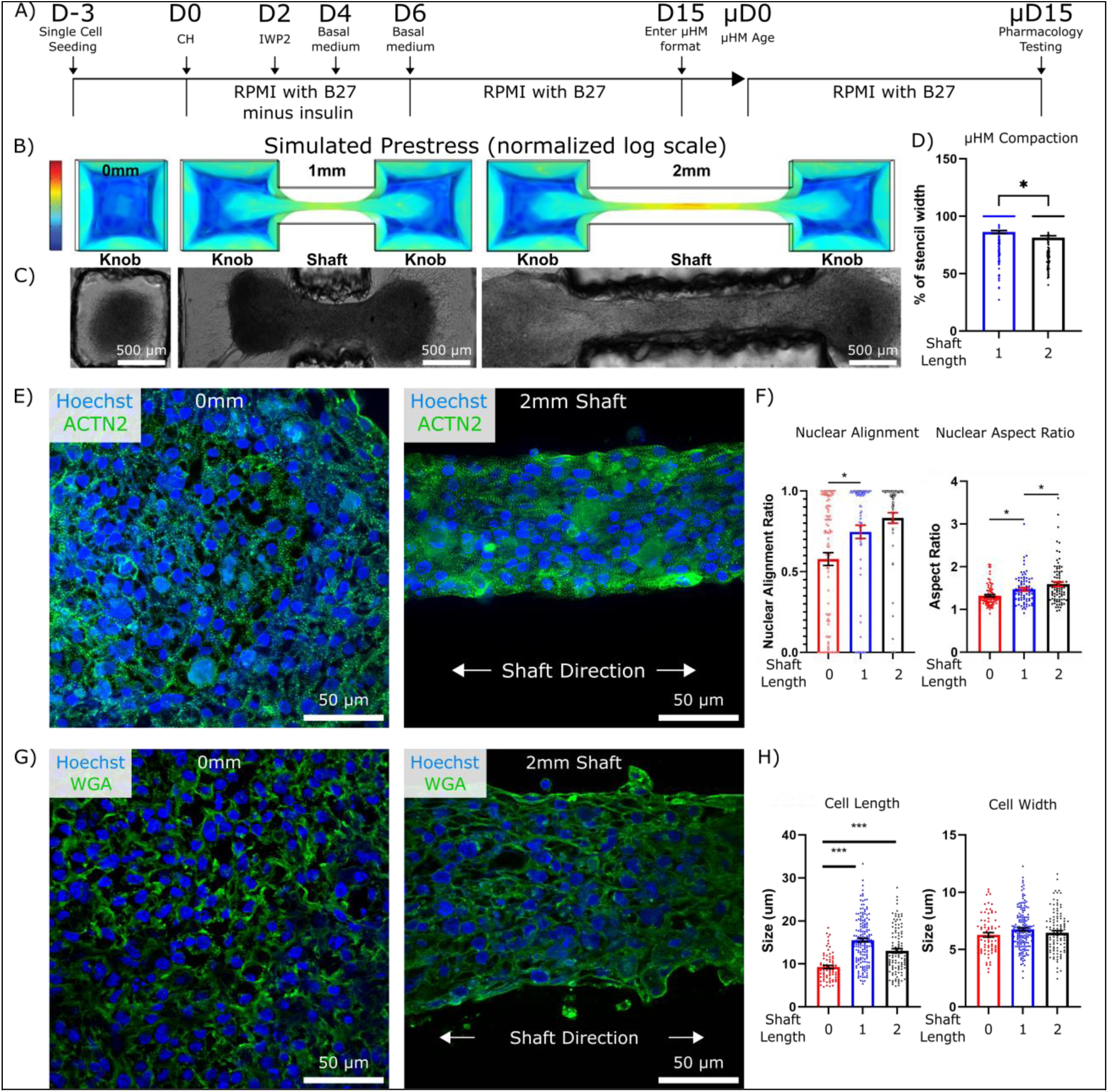
Prestress Regulates Cell Morphology In Micro-Heart Muscle (µHM). **A**) Timeline of small molecule-based differentiation of iPSC to cardiomyocytes, the subsequent seeding into the µHM format, and the measurements of cell morphology, electrophysiology, and pharmacology experiments at the terminal time point. **B)** COMSOL modeling images depicting increased tissue preload with increased shaft length. **C)** Representative bright field images taken at µHM day 15 (µD15) showing grossly similar tissue morphology regardless of tissue size. **D)** Quantification of µHM shaft lateral compaction at µD15 (*n* > 150 µHM). **E)** Representative confocal images of nuclei (Hoechst/blue) and sarcomeric α–actinin (green) of 0mm and 2mm tissues at µD15. **F)** Quantification of the effects of tissue geometry on nuclear alignment and nuclear aspect ratio at µD15 (*n* > 70 nuclei from 6-8 µHM per size). **G)** Representative confocal images of nuclei (Hoechst/blue) and cell membrane (wheat germ agglutinin/green) of 0mm and 2mm tissues at µD15. **H)** Quantification of the effects of tissue geometry on cell length and width at µD15 (*n* >100 cells from 6-8 µHM). *: p < 0.05, **: p < 0.005, ***: p < 0.0005, error bars: *SEM*)

Despite the overall similarities in gross morphology between μHM with 1mm vs. 2mm shaft lengths, staining for sarcomeric α-actinin and cell membrane in longitudinal cryosections of µHM suggested that tissue geometry had marked effects on cardiomyocyte structure (Fig. 1E, G). Nuclear alignment and aspect ratio, especially useful metric of cell organization in highly dense 3D tissues, both increased with increased μHM shaft (Fig. 1F). Cell length also increased as μHM shaft length increased from 0mm to elongated (1 & 2mm tissues) geometries (Fig. 1H). Interestingly, although cell length increased with shaft length, cell width was unchanged (Fig. 1H). This is consistent with eccentric hypertrophy, which is observed *in-vivo* in the context of increased preload.^34^ Consistent with model predictions, changes to nuclear and cell morphology were most pronounced in the elongated “shaft” region of μHM (Fig. S1). Collectively, these results confirm that increasing µHM length induces increasing levels of tissue prestress, which then directly modulates cardiomyocyte morphology and alignment.

### Action Potential and Calcium Handling Kinetics are Modulated by Tissue Prestress

We next investigated potential effects of prestress on electrophysiology of spontaneously beating µHM through optocardiography. Broadly, whole-tissue action potential and calcium transient waveforms displayed marked remodeling as μHM shaft length increased (Fig. 2A). Quantitative analysis of several key parameters describing the kinetics of action potentials and calcium transients (e.g. time to 90% action potential duration, APD_90_, and the time required for calcium transients to decay from peak to 25% of peak amplitude, τ_75_) confirmed the robustness of these prestress-induced global changes across a large number of μHM amassed from multiple differentiation batches (Fig. 2B, S2C). Importantly, analysis of individual action potential and calcium transient kinetic parameters suggested no measurable changes in timing whether the entire tissue or a smaller sub-region was analyzed (Fig. S2A). This suggests that µHM electrophysiology is dominated by the regions of highest prestress within individual tissues. Consistent with a more mature phenotype,^2, 26^ µHM with longer shafts exhibited lower spontaneous beating rates (Fig. 2C). Beat-rate corrected action potential upstroke duration (cUPD) and APD_90_ also increased with prestress (Fig. 2B, C, S2C).

**Figure 2.**
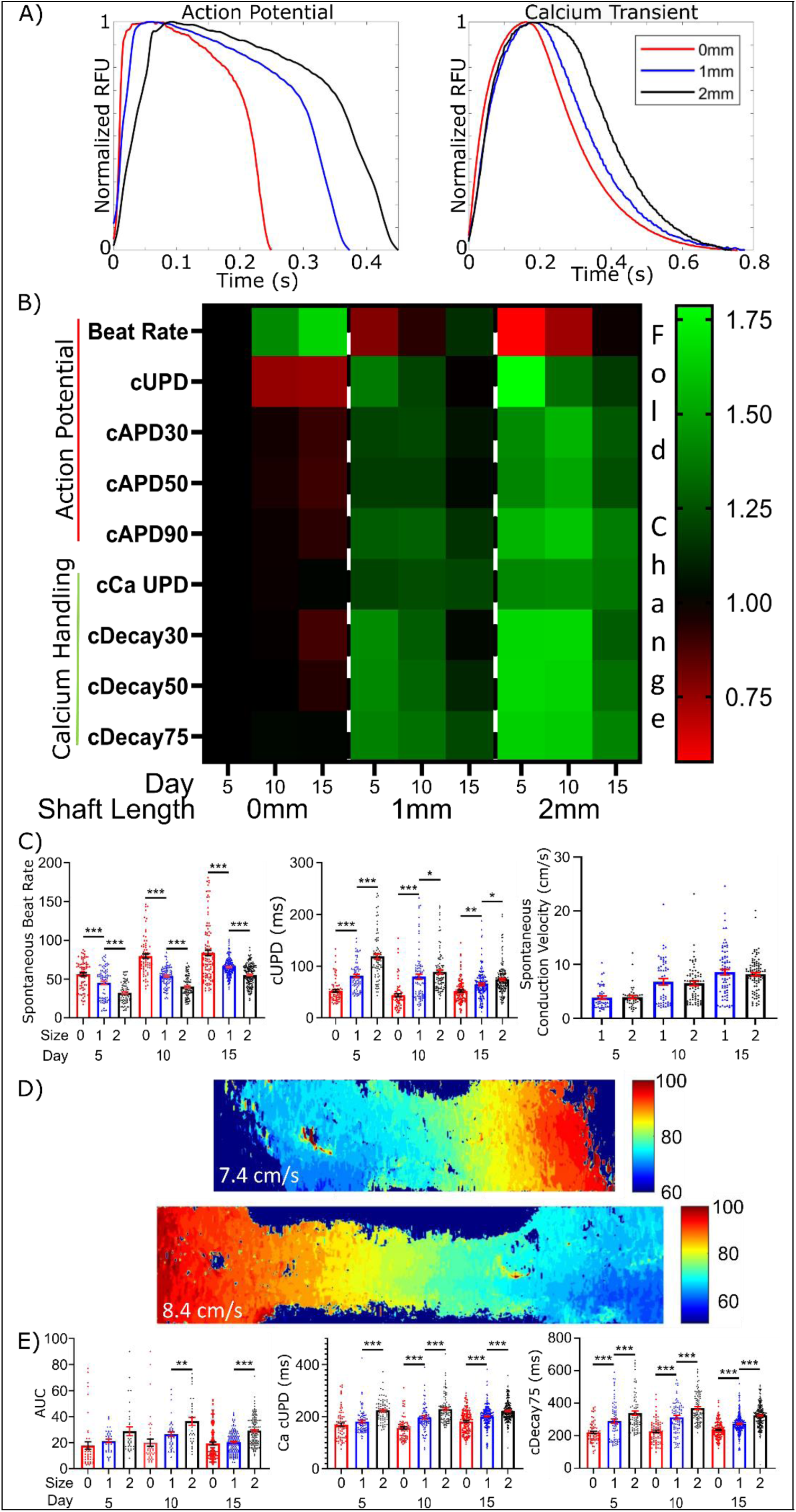
Tissue Prestress Regulates µHM Electrophysiology. **A)** Representative action potential and calcium transient waveforms for 0mm, 1mm, and 2mm µHM at µD15. **B)** Electrophysiology heatmap of action potential and calcium handling dynamics for µHM of shaft length 0, 1, & 2mm long. Data is normalized to µD5 0mm data for given parameter. **C)** Select data contained within the heatmap shown in panel B: spontaneous beat rate, beat-rate corrected action potential upstroke duration, and spontaneous conduction velocity. **D)** Representative activation maps of 1mm and 2mm µHM at µD15. **E)** Area under the curve of calcium transient waveforms, beat-rate corrected calcium transient upstroke duration, and beat-rate corrected calcium transient decay 75 for 0, 1, & 2mm µHM up to µD15 (*n* > 100 µHM, *: p < 0.05, **: p < 0.005, ***: p < 0.0005, error bars: *SEM*).

Increases in action potential upstroke duration are most likely to be reflective of one of three possibilities: 1) decreased sodium current, which would suggest less mature tissues,^37^ 2) increased overall influx of ions into the cardiomyocytes during depolarization,^38^ or 3) increased gap-junction coupling between cells, which slows action potential and calcium upstroke timing in tissues.^39^ To address these distinct possibilities, we first analyzed the spontaneous conduction speed across µHM, which is partially controlled by both sodium channel function and gap-junction coupling. Although distinctive in their overall action potential morphology, we observed no significant difference in conduction speed between 1 and 2mm µHM at any time point.

However, both geometries exhibited conduction speed increased over time (Fig. 2C, D).

In contrast to conduction speed, which was consistent between μHM with 1 and 2mm shaft lengths, we observed marked differences in μHM calcium handling linked with shaft length. Both the upstroke duration and decay times of the calcium transients increased with increasing tissue length and prestress (Fig. 2E). Together, these changes led to an increase in the total amount of calcium influx into the cardiomyocytes, as measured by the area under the curve (AUC) of calcium transient waveforms (Fig. 2E). Altogether, these results are most consistent with increases in total ion influx, rather than enhanced gap junction function or sodium channel impairment, as the predominant mechanism for the observed increase in APD and upstroke timing as a function of tissue prestress.

### A Tissue Prestress Threshold is Necessary for Development of Functional Sodium Channel

We next used a series of drugs selected to block unique ion channels to determine what channels were most affected by tissue prestress to cause the observed electrical remodeling. We first assessed the effects of tissue prestress on the Nav1.5 sodium channel, responsible for the action potential upstroke in the adult human cardiomyocyte and partially for the upstroke in matured iPSC-cardiomyocytes,^40^ through the use of saxitoxin. Strikingly, aligned tissues (µHM with either 1 or 2mm shaft length) exhibited an appropriate dose-response with increasing cUPD as *I_Na_* block increased (Fig. 3A, B), whereas tissues with no prestress or preferential alignment (0mm) exhibited no such response (Fig. 3A, B). These observations suggest that a prestress threshold is necessary for Nav1.5 ion channels to make functional contributions to the action potential. Moreover, although the 1 & 2mm tissues show similar *I_Na_* blockage curves in relation to cUPD, μHM with 2mm shaft length consistently maintain a higher cUPD. These observations strongly suggest that the increased upstroke time in µHM with anisotropic, elongated geometry compared to the 0mm µHM (Fig. 2A,B, 3B) is due to the prevalence of increased *I_Na_*, as opposed to a Nav1.5 deficiency. This would lead to a greater influx of Na^+^, and subsequently Ca^2+^, into the cells and prolong the upstroke duration of the action potential and calcium transient.^38^ This possibility was corroborated by a decrease in spontaneous conduction velocity in the presence of saxitoxin (Fig. S3A).

**Figure 3.**
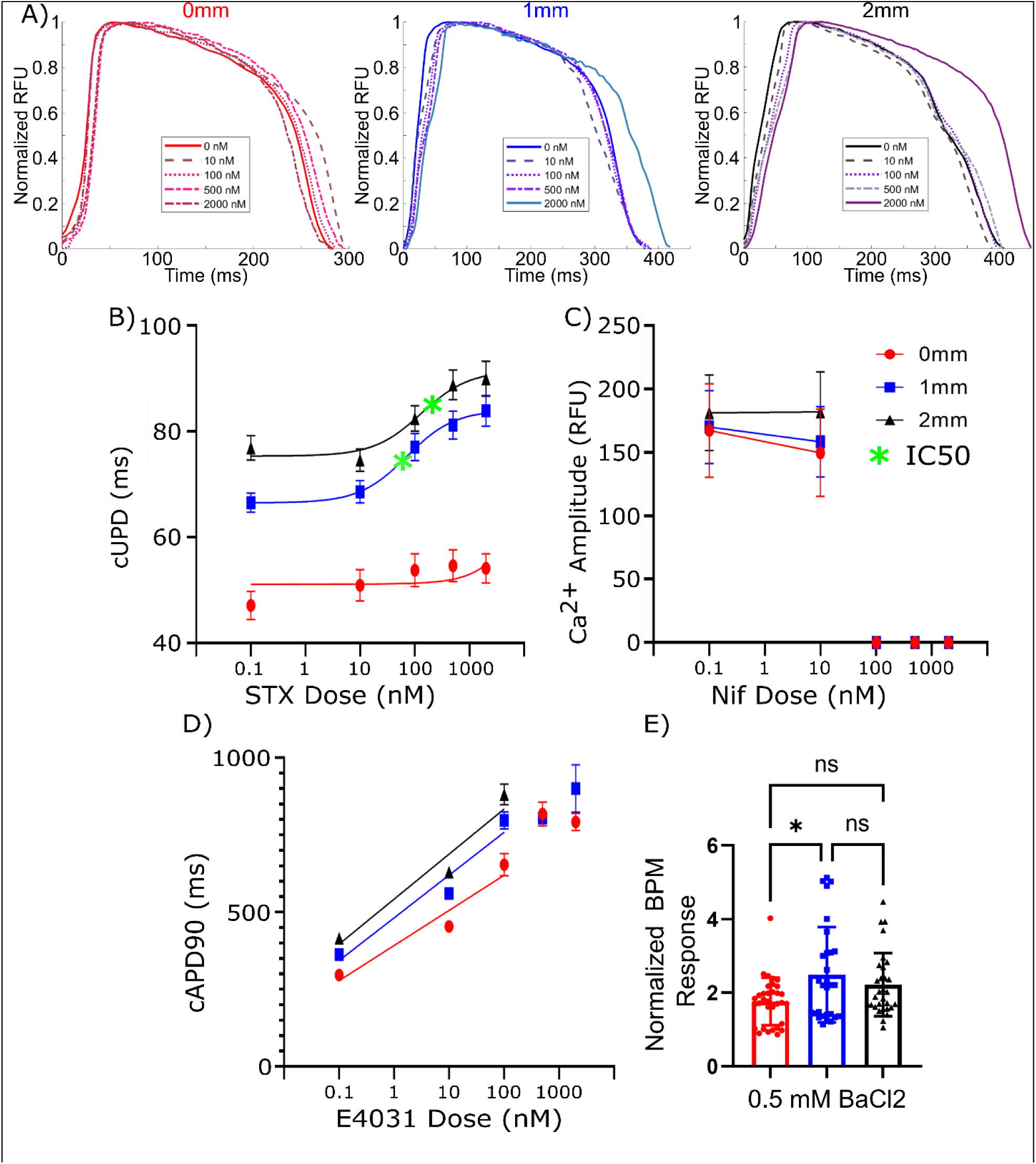
Pharmacologic Inhibition of Ion Channels. **A**) Representative action potential waveforms for 0mm, 1mm, and 2mm µHM after sequential saxitoxin doses. **B-F)** Response of µHM to pharmacologic inhibition of ion channels. Effect of **B)** saxitoxin on upstroke duration, **C)** nifedipine on calcium transient amplitude, **D)** E4031 on APD_90_, and **E)** 0.5 mM BaCl_2_ on spontaneous beat rate. (*n* = 20-30 µHM, *: p < 0.05, **: p < 0.005, error bars: *SEM*)

In contrast to observations linking prestress to functional sodium currents, blockade of either the *L*-type calcium current (*I_CaL_*) with nifedipine or the rapidly rectifying potassium current (*I_Kr_*) with E4031 revealed no marked prestress-dependent changes (Fig. 3C, D, S3B, C). This suggests that the calcium transient shifts stemming from μHM geometry changes (Fig. 2A, B) are most likely due to increases in *I_Na_* that allow more calcium to enter the cell, whereas calcium channels themselves are not significantly affected by tissue prestress. Likewise, increases in APD_90_ with tissue prestress are unlikely to be caused by changes in *I_Kr_*, and instead are more likely to be caused by increased ion influx (Fig 3.2D). Although the elongated µHM gain functional *I_Na_*, *I_CaL_* remains necessary for action potential initiation, as observed by the block of AP altogether at high dose of nifedipine, and the finding that only a small component of conduction velocity is directly linked to *I_Na_* (Fig. S3A). These findings highlight that at this timepoint, these tissues remain somewhat immature even with the highest level of tissue prestress.

Based on observations of changes in spontaneous tissue beating rate and increased sodium channel function, we next used pharmacologic approaches to address functional changes to *I_K1_*, which regulates resting membrane potential. To determine if this current increases alongside *I_Na_*, *I_K1_* was blocked with a high dose of BaCl_2_. While all tissue geometries responded to BaCl_2_ by increasing their spontaneous beat rate, we observed a smaller effect on spontaneous beat rate and APD_90_ changes in 0mm as compared to elongated tissues (Fig. 3E, S3D). It is thus possible that the lower spontaneous beat rate with increasing tissue length (Fig. 2C) is due to increased expression of Kir2.1, the major protein linked to *I_K1_*. This protein has been shown to be localize to the cell membrane in a focal adhesion dependent manner, which may explain tensional regulation.^41^ Additionally, and consistent with our findings that *I_Na_* and *I_K1_* are the most likely currents to be regulated by tissue prestress, the main proteins encoding these currents, Kir2.1 and Nav1.5 have also been shown to co-traffic^42^ and form complexes within cardiomyocytes.^43^

### Potential Molecular Mechanisms of Prestress Regulation of Ion Channel Expression

To identify potential molecular mechanisms causing differential *I_Na_* in µHM, we first quantified RNA expression for a variety of structural proteins and action potential related ion channels and calcium handling proteins. Although we observed significant upregulation of several transcripts encoding structural proteins when comparing µHM to input iPSC-CM (Fig. S34), differences in expression of these same transcripts as a function of μHM geometry was negligible (Fig. 4A). This is consistent with previous work reporting an apparent decoupling between transcript levels and function of individual ion channels.^44^ Quantification of cardiomyocyte entry into proliferative stages of the cell cycle via Ki67 staining likewise did not indicate an effect of µHM geometry (Fig. S5). Altogether, these data suggest that prestress induced changes observed in electrophysiology are not due to large shifts in ion channel expression or changes in the cell cycle that would be expected if the cause of electrical remodeling were a global change in cardiomyocytes maturation state.

**Figure 4.**
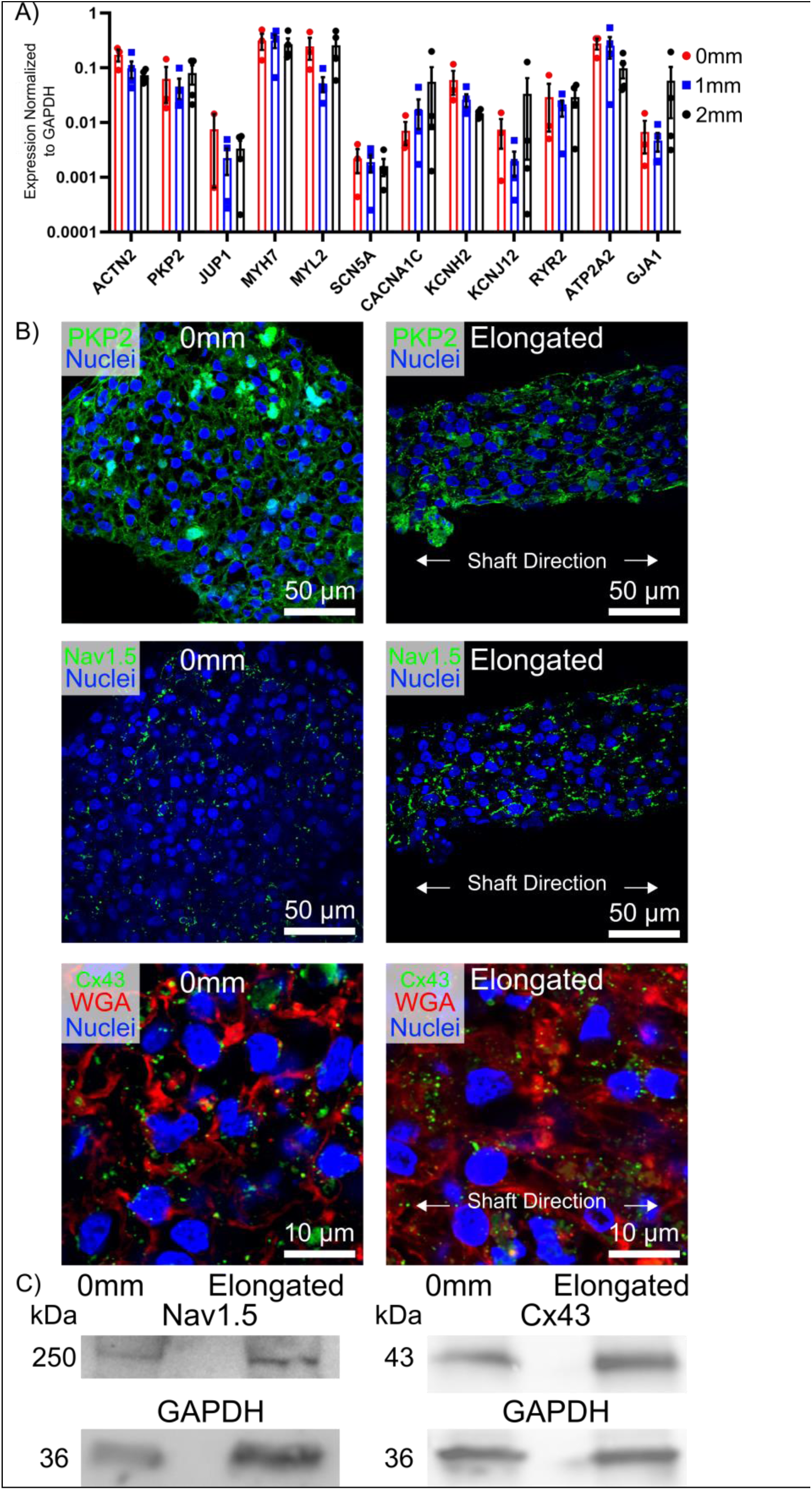
Nav1.5 and Gap Junctions are Regulated Post-Translationally. **A**) Relative expression of 12 genes encoding for mechanical and action potential-related ion channels in µHM at µD15: ACTN2, PKP2, JUP1, MYH7, MYL2, SCN5A, CACNA1C, KCNH1, KCNJ12, RYR2, ATP2A2, and GJA1. Error bars: *SEM*. Each data point represents RNA from pooled µHM from a single independent iPS-CM differentiation batch (*n* = 4). **B)** Representative staining of 0mm and elongated µHM at µD15 for: Plakophilin-2, Nav.15, and Connexin-43. **C)** Representative Western blot of 0mm and elongated µHM at µD15 for: Nav1.5, Connexin-43, and GAPDH.

As ion channel function is strongly linked to both protein expression and localization, we next assessed the expression and localization of key membrane-localized proteins in µHM. Plakophilin-2, a major component of the desmosome associated with the function of both Nav.5 and connexin-43 (the main gap junction protein in cardiomyocytes),^45^ was localized correctly to cell-cell junctions regardless of tissue length and prestress (Fig. 4B). In contrast, we observed a trend toward increased Nav1.5 and connexin-43 (Cx43) expression in the shaft region of elongated tissues, without any indication of mis-localization away from the cell membrane in 0mm tissue (Fig. 4B).

Analysis of bulk protein levels within tissues suggested a trend toward increased Cx43 protein expression in elongated µHM (1 and 2mm shafts) (Fig. 4C, S6), similar to observations that geometries expected to induce higher prestress enhanced Cx43 expression in other studies.^21^ However, the trend we observed was not statistically significant (*p =* 0.099, 2-way *t*-test over 5 batches of µHM). Moreover, analysis of overall Na_V_1.5 protein levels did not reveal any similar trend of upregulation (Fig. 4C). Altogether, these results suggest that post-transcriptional, if not post-translational mechanisms (e.g., trafficking) link changes in tissue prestress to changes in sodium channel function in μHM.

### Generation of Plakophilin 2 Knockout Micro-Heart Muscle Arrays

Cardiomyocytes contain specialized mechanical junctions called desmosomes,^46^ which enable cell-cell connections within the high stress environment of the heart, and are comprised of: desmoglein-2, desmocollin-2, plakoglobin, plakophilin-2, and desmoplakin. Consistent with the hypothesis that desmosomes are critical for prestress sensing, patients with genetic mutations for desmosome proteins, most commonly plakophilin-2, show increased sensitivity to elevated mechanical loading on the heart.^47–52^ Further, physical activity has been shown to increase the risk of sudden cardiac death by five-fold in these patients.^53^ Importantly, many ion channels related to the action potential preferentially localize to the desmosome,^54^ and the desmosome is linked to the expression of proteins such as Na_V_1.5 and Cx43.^55^ These observations show that cell-cell junctions play a crucial role in how cardiomyocytes respond to both normal and pathologic heart development. To determine whether plakophilin-2 plays a role in regulating iPS-CM electrical remodeling in response to tissue prestress, isogenic, PKP2^-/-^ iPSC were generated using CRISPR/Cas9. The absence of PKP2 protein in differentiated iPS-CM was confirmed via immunoblotting (Fig. 5A, S7).

**Figure 5.**
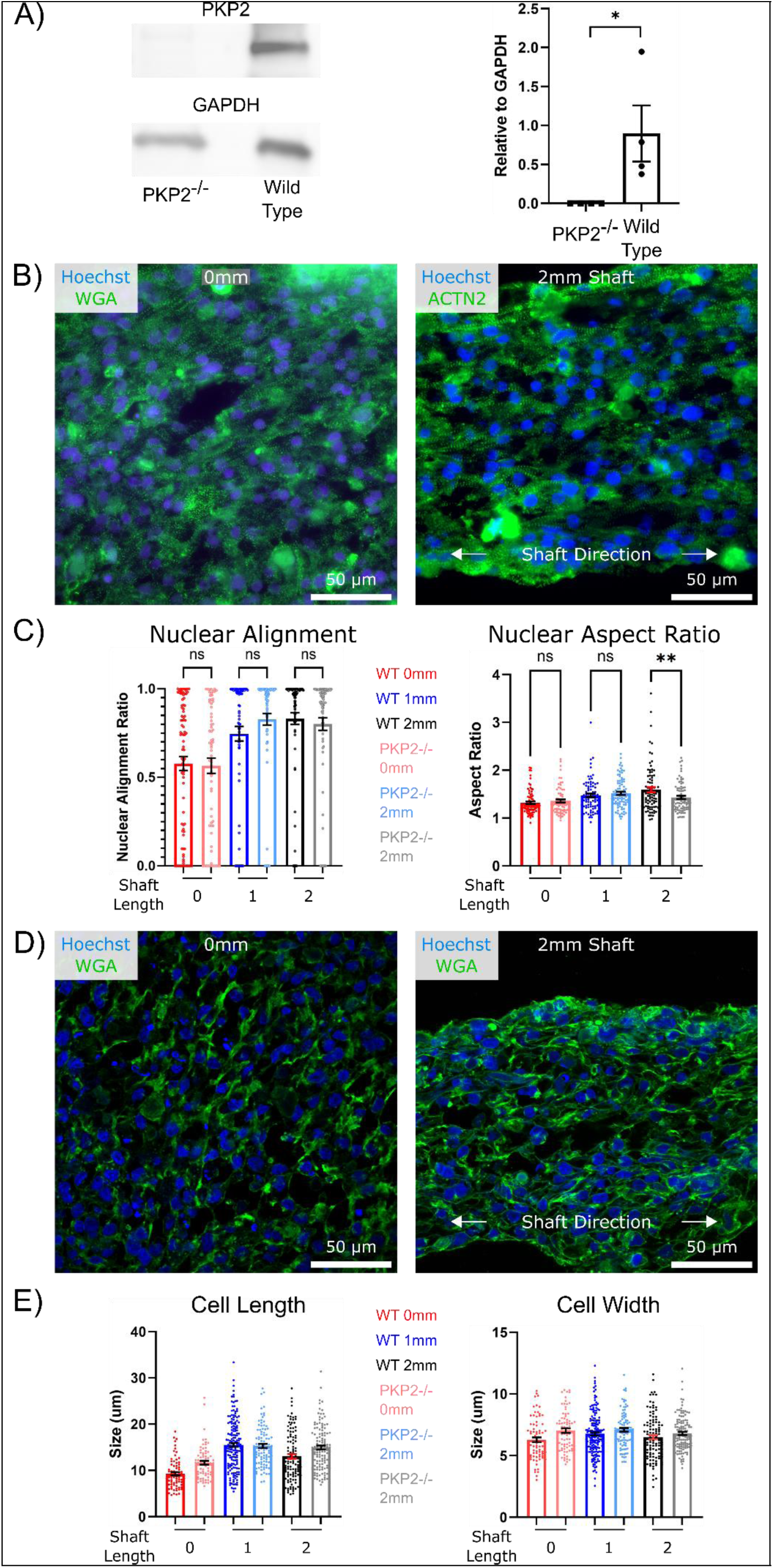
Prestress Regulates Cell Morphology In PKP2^-/-^ µHM. **A**) Representative images and quantification of western blot for PKP2 and GAPDH in PKP2^-/-^ and isogenic control iPS-CM. (*n* = 4 differentiation batches) **B**) Representative images of nuclei (Hoechst/blue) and sarcomeric α–actinin (green) of PKP2^-/-^ 0mm and 2mm PKP2^-/-^ µHM at µD15. **C)** Quantification of the effects of tissue geometry on nuclear alignment and nuclear aspect ratio at µD15 in wild type and PKP2^-/-^ µHM (*n* > 70 nuclei from 6-8 µHM per size). **D)** Representative images of nuclei (Hoechst/blue) and cell membrane (wheat germ agglutinin/green) of 0mm and 2mm PKP2^-/-^ µHM at µD15. **E)** Quantification of the effects of tissue geometry on cell length and width at µD15 in wild type and PKP2^-/-^ µHM (*n* >100 cells from 6-8 µHM, *: p < 0.05, **: p < 0.005, ***: p < 0.0005, error bars: *SEM*).

### Loss of Plakophilin-2 Does Not Affect Cell Morphology within µHM

Analysis of sarcomeric α-actinin and overall cell membranes in 0mm and 2mm PKP2^-/-^ µHM (Fig. 5B, D) did not reveal any gross abnormalities. Likewise, quantification of nuclear alignment suggested no significant differences in overall cell alignment between control and PKP2^-/-^ µHM. A similar trend was observed with respect to nuclear aspect ratio, although the absolute level of nuclear anisotropy was slightly lower in 2mm PKP2^-/-^ µHM compared to 2mm controls (Fig. 5C). The correlation of increased tissue and cell length was also maintained, along with the lack of change in cell width (Fig. 5E). Consistent with COMSOL model predictions (Fig. 1B) and results from control tissues, changes to nuclear and cell morphology were most pronounced in the tissue shaft (Fig. S8). These observations are consistent with findings that PKP2 null mouse embryos are capable of forming the primitive heart tube.^56^ While likely incapable of withstanding the prestress levels present in postnatal or even late stage embryonic hearts, the PKP2 knockout cardiomyocytes generated here are still likely to retain other protein systems critical for cell-cell mechanical coupling, including cadherins and catenins.^57^ These alternative protein systems are likely sufficient to induce prestress-regulated morphology changes in these tissues despite the loss of PKP2 and the potential impacts of this loss on nascent desmosome structure and mechanical integrity.

### PKP2 Knockout μHM Exhibit Blunted Electrical Remodeling in Response to Prestress

We next investigated the role that PKP2 plays in the prestress-regulated electrophysiology changes we observed in control μHM. Broadly, although we observed some differences in the action potential and calcium transient waveforms as a function of μHM geometry in PKP2^-/-^ µHM (Fig. 6A), prestress-induced changes in overall waveform morphology were more subtle than in controls. Unlike control tissues, those without PKP2 exhibited no differences in spontaneous beat rate across different μHM shaft lengths µHM day 15 (Fig. 6B). Likewise, action potential upstroke duration was largely insensitive to prestress (Fig. 6B, S9). By day 15 of culture within μHM (µD15), both elongated (1 & 2mm) tissues maintained higher upstroke durations compared to the 0mm tissues, but not from one another.

**Figure 6.**
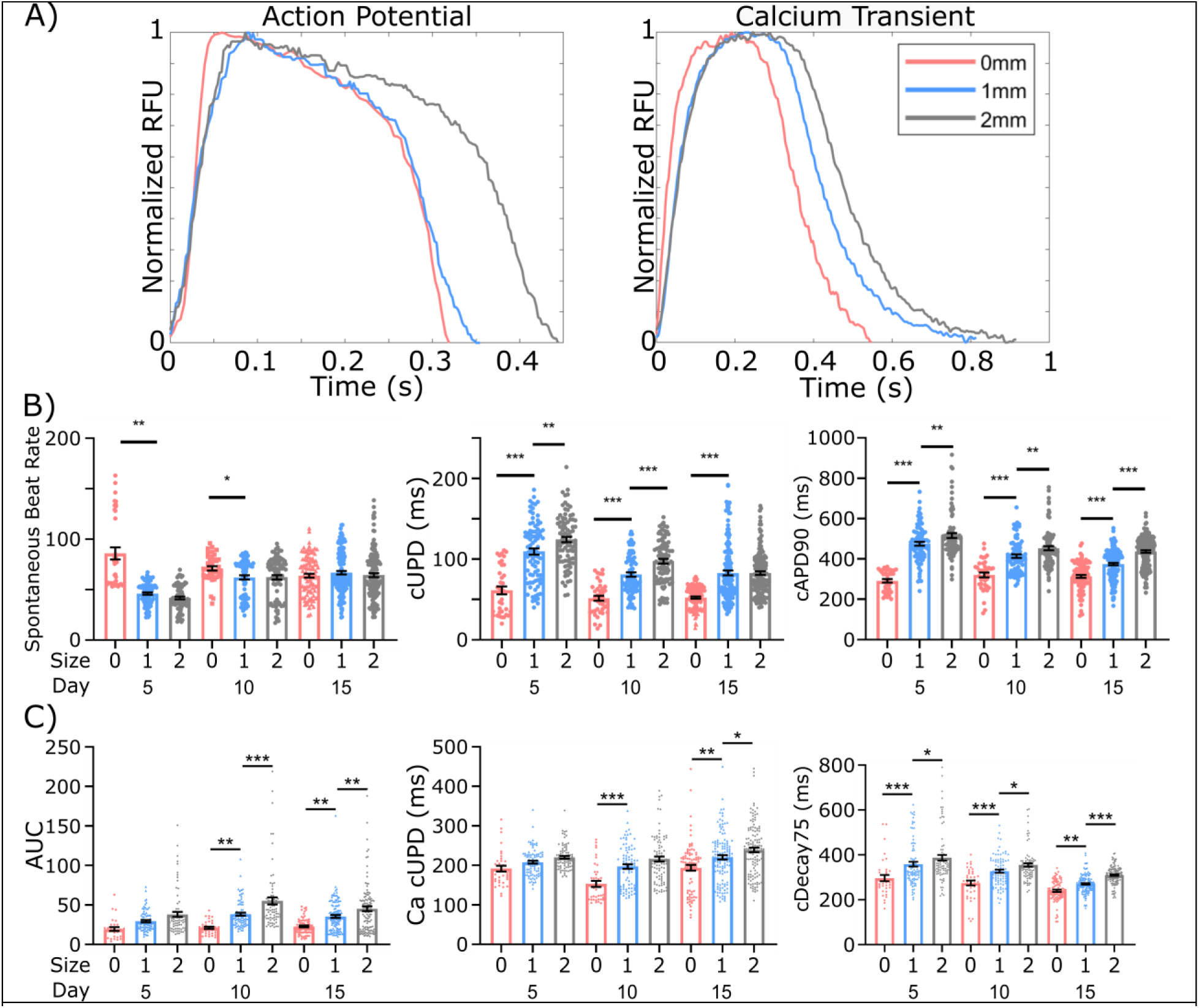
Loss of PKP2 Mutes Tissue Prestress Regulation of µHM Electrophysiology. **A**) Representative action potential and calcium transient waveforms for 0mm, 1mm, and 2mm PKP2^-/-^ µHM at µD15. **B)** Spontaneous beat rate, beat-rate corrected action potential upstroke duration, and beat-rate corrected APD90, and **C)** area under the curve of calcium transient waveforms, beat-rate corrected calcium transient upstroke duration, and beat-rate corrected calcium transient decay 75 for 0, 1, & 2mm PKP2^-/-^ µHM, up to µD15 (*n* = 40-100 µHM, *: p < 0.05, **: p < 0.005, ***: p < 0.0005, error bars: *SEM*).

Despite a general lack of changes in action potential waveforms, calcium handling differences across μHM geometries appeared to be conserved in the absence of PKP2, with the area under the curve, Ca^2+^ transient upstroke duration and decay_75_ all differentially regulated by μHM geometry at µD15 (Fig. 6C).

### PKP2 Is Necessary For Prestress Regulation Of Sodium Current

To determine if prestress regulation of sodium current is desmosome dependent, we again used pharmacologic inhibition of ion channels to view their contributions to the action potential and calcium handling. Unlike control μHM, which exhibited marked action potential morphology changes when *I_Na_* was blocked with saxitoxin, PKP2^-/-^ µHM exhibited no significant action potential waveform changes in response to this drug, regardless of tissue geometry (Fig. 7A,B, S10A).

**Figure 7.**
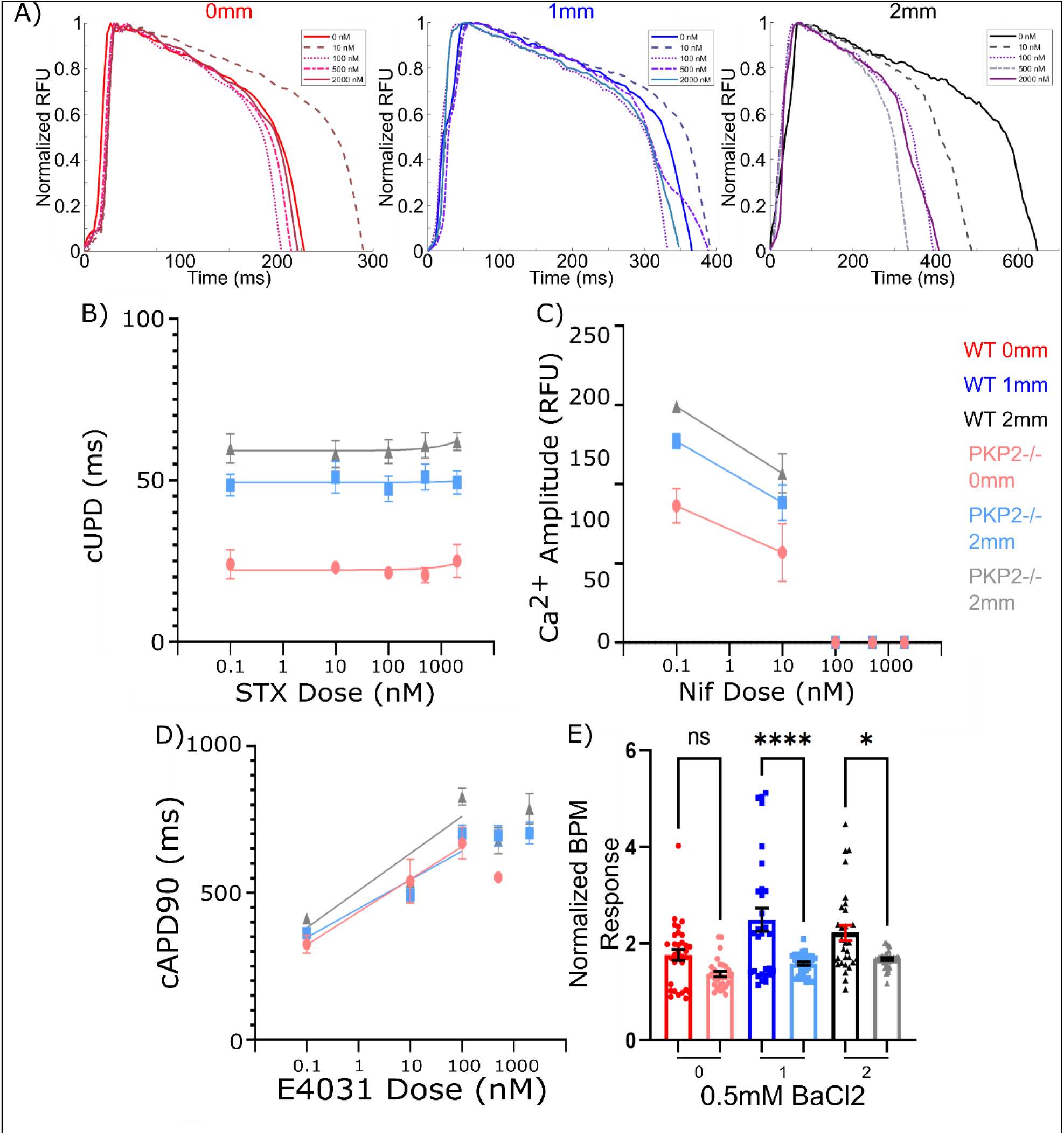
Pharmacologic Inhibition of Ion Channels in PKP2^-/-^ μHM. **A**) Representative action potential waveforms for 0mm, 1mm, and 2mm PKP2^-/-^ µHM after sequential saxitoxin doses. **B-F)** Response of PKP2^-/-^ µHM to pharmacologic inhibition of ion channels. Effect of **B)** saxitoxin on upstroke duration, **C)** nifedipine on calcium transient amplitude, **D)** E4031 on APD_90_, and **E)** 0.5 mM BaCl_2_ on spontaneous beat rate. (*n* = 6-24 µHM). Error bars: *SEM*

Blockage of Cav1.2 with nifedipine had a similar effects in PKP2 null μHM as in controls, with a 100 nM dose being sufficient to block action potential initiation (Fig. 7C). Strikingly, however, while 10 nM of nifedipine had minimal effects on the amplitude of the calcium transients for control μHM (Fig. 3.3C), this dose markedly decreased Ca^2+^ transient amplitude in PKP2^-/-^ tissues. The slope of this decrease appears relatively similar for all tissue sizes, suggesting similar Cav1.2 kinetics. Likewise, while 10 nM of nifedipine led to no difference in the upstroke duration of the calcium transient in control μHM (Fig. S3B), this led to a significant decrease in PKP2^-/-^ tissues (Fig. S10B). These results suggest the higher action potential cUPD for the 2mm PKP2^-/-^ tissues is due to increased calcium influx through L-type channels, as *I_Na_* makes no contribution to the action potential upstroke in these PKP2 deficient tissues.

Blockage of hERG with E4031 showed a nearly identical effect on the cAPD_90_ of PKP2^-/-^ μHM as was observed in control tissues (Fig. 3D), with doses of 500 nM and above inhibiting the ability for cardiomyocytes to repolarize (Fig 7D). Additionally, the slope of the sensitivity across all tissues appears the same, suggesting minimal differences in hERG activity across μHM shaft lengths. However, a dose of 100 nM significantly decreased the spontaneous beat rate in PKP2^-/-^ µHM (Fig. S10C), whereas the effect of this dose on WT µHM was minimal (Fig. S3C). A similar effect was seen on the action potential amplitude as well. Thus, although there does seem to be a degree of prestress regulation of calcium handling and action potential duration in PKP2^-/-^ µHM, the degree of this regulation appears to be significantly less than in isogenic controls, and appears to be driven by changes in activity of hERG and *L*-type calcium channels.

Finally, we examined potential changes in *I_K1_* in PKP2 deficient μHM. In contrast to control μHM, PKP2^-/-^μHM of all shaft lengths exhibited minimal differences in the spontaneous beat rate in the presence of a blocking dose of BaCl_2_ (Fig. 7E). This response correlates with the lack of a prestress regulated spontaneous beat rate response seen in these tissues (Fig. 6B). We further assessed the expression and localization of Nav1.5 and Cx43 in elongated PKP2^-/-^ µHM by immunostaining. Although both of these proteins were present, overall levels appeared to be markedly decreased compared to control μHM (Fig. 8A). The majority of cells (> 90%) did not stain for either of these proteins. Analyzing bulk protein levels over entire µHM indicated a potential trend toward increased Cx43 protein expression in elongated µHM (1 and 2mm shafts; Fig. 8A, S11). As was observed in isogenic control μHM, however (Fig. 4C, S6), this trend was not statistically significant. Western blotting for overall Na_V_1.5 protein levels revealed that the protein is likely present, but at a substantially lower level than in isogenic control μHM (Fig. 8). Altogether, these results suggest that while cardiomyocytes are capable of adapting morphologically to mechanical prestress without PKP2, this protein is necessary for proper electrophysiologic adaptation and ion channel expression.

**Figure 8.**
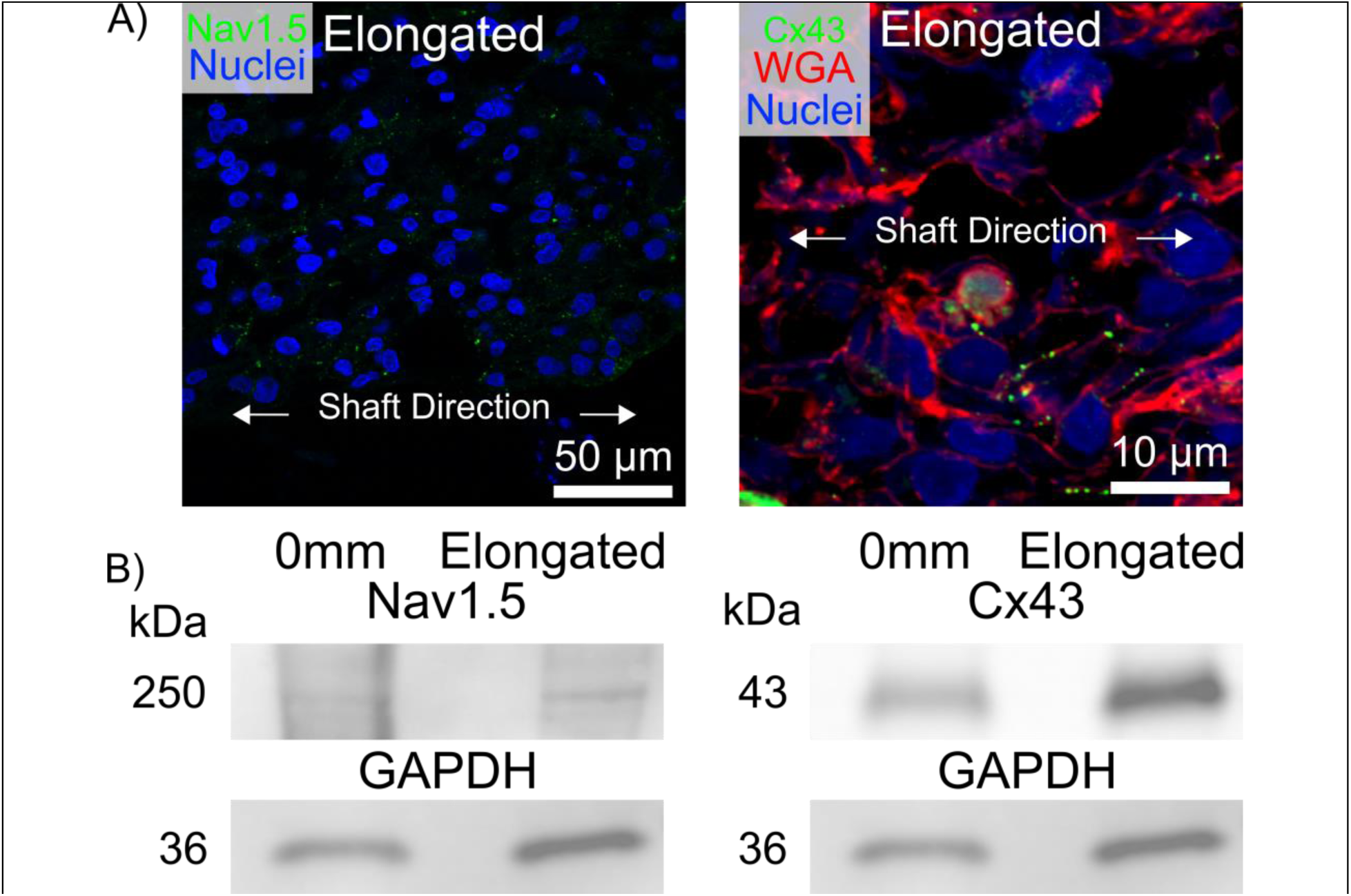
Nav1.5 and Cx43 Expression Are Plakophilin-2 Dependent. **A**) Representative staining of 0mm and elongated PKP2^-/-^ µHM at µD15 for Nav.15, and Connexin-43. B) Representative Western blot of 0mm and elongated µHM at µD15 for: Nav1.5, Connexin-43, and GAPDH.

## Conclusions

Human induced pluripotent stem cell-derived cardiomyocytes show promise in predicting drug-induced arrhythmias more accuracy compared to alternative *in-vitro* assays.^1^ Tissue engineering approaches have been applied to enhance iPSC-CM electrophysiologic maturity.^2, 14, 16, 20, 21, 31, 58^ To test the hypothesis that prestress on cardiomyocytes in the 3D setting of engineered tissues plays a role in electrical remodeling, we tested the impact of micro-tissue geometry on action potential and calcium transients. The results seen reveal that tissue prestress regulates activity of specific ion channels to drive remodeling of the cardiac action potential. Importantly, prestress alone is sufficient to drive a lower spontaneous beat-rate and form functional Na_V_1.5 current after only two weeks of culture. Prestress also appeared to regulate the expression of proteins, including Kir2.1 and Cx43, although to a lesser degree. However, no level of prestress was able to eliminate spontaneous beating at this timepoint, and action potential upstroke is still highly Ca^2+^ dependent. These observations suggest that applying additional exogenous stretch and/or combining geometric constraints with chemical and chemical stimuli may be required to induce further electrophysiological maturation.^26, 59^

The observations seen regarding Cx43 and Nav1.5 expression suggest that robust protein expression differences may require longer culture time in the µHM environment. Given a half-life of 1-3 hours^60^ and a disconnect between gene and protein expression^44^ for Cx43, it is likely that changes we observe in µHM upstroke duration (Fig. 3B) are related to post-translational modifications of Nav1.5, and potentially Cx43. Changes in trafficking may be a key post-translational change in Cx43 and Nav1.5, which has been previously shown to be dependent on tissue stress and related cytoskeletal structures (microtubules and focal adhesions).^41, 60–63^ These trafficking changes may be sufficient to drive marked functional changes in *I_Na_* without requiring gross changes in the overall levels of the transcript encoding for the functional pore-forming subunit of the sodium channel. Separately, proper trafficking of Cx43 is dependent on the GJA1-20k isoform.^64^ A similar mechanism exists for Nav1.5, in which the β2 subunit regulates trafficking to the cell membrane.^65^ It remains unknown if and how these and other ion channel isoforms are regulated by tissue prestress, and how they may contribute to the prestress-regulated electrophysiology changes seen. A small increase in BaCl_2_ sensitivity was seen with increasing tissue length, suggesting Kir2.1 expression increases alongside Nav1.5 expression, in concurrence with previous studies.^42, 43^ Changes seen in µHM spontaneous beat rate and sensitivity to BaCl_2_ suggest the resting membrane potential of these tissues may be altered by prestress, which could affect the opening and activation kinetics of Nav1.5.^66^

Ultimately, the data shown here reveals that a prestress threshold is necessary for sodium channel function in cardiomyocytes, and that changes in electrophysiology seen between engineered heart tissues and iPS-CM are due to changes in specific ion channels and are unlikely linked to global shifts in maturation. These observations suggest the cardiomyocytes adapt to increased levels of tissue prestress by increasing their calcium influx and handling, which is achieved by increasing the sodium current to increase the ion influx into the cell and prolongs the action potential.

To understand the mechanisms by which tissue prestress regulates developmental ion channel expressionPKP2^-/-^ µHM of multiple geometries were created. Although strong differences in electrophysiology were seen between PKP2^-/-^ and isogenic control µHM and their response to varying levels of tissue prestress, PKP2^-/-^ µHM did not exhibit marked arrhythmias under the present experimental conditions. It is potentially the case that higher levels of mechanical loading, adrenergic stress and/or further maturation are necessary to observe arrhythmias seen commonly in patients with PKP2 mutations.^67^ The results seen reveal that although the loss of PKP2 does not completely inhibit prestress-induced morphology changes on the cardiomyocytes, it does starkly inhibit the electrophysiology changes seen in wild-type tissues due to tissue geometry and prestress, and may lead to an over-reliance on calcium intake to respond to changes. These changes are due to inhibitions in the expression of a variety of ion channels. There was no functional Nav1.5 expression in plakophilin-2 deficient tissues. Cav1.2, hERG, Kir2.1, and Cx43 also appeared to be dependent as well, although to a lesser degree. Ultimately, the data shown here reveals that plakophilin-2 is necessary for cardiomyocytes to properly adjust their physiology according to tissue prestress.

The work described here details the specificity with which prestress regulates certain aspects of cell electrophysiology, while others are not dependent on this type of mechanical loading. This understanding of cardiomyocyte development will allow for the creation of future *in-vitro* models that more accurately model adult human cardiac response, and can be used to study disease pathology, predict cardiac drug toxicity, and study heart development.

Future work can study the role increased levels of prestress, and potentially dynamically loading stress, affect cardiomyocyte physiology.^13, 59^ Incorporating this work with studies utilizing differential levels of afterload^23^ to determine how they independently regulate electrophsyiology would also be of interest. Previously published studies utilizing other engineered heart tissue systems that incorporate electrical pacing has seen the formation of t-tubules and higher levels of tissue conduction velocity.^20, 21^ Other work has utilized tissue designs that more accurately mimic the type of loading cardiomyocytes are subjected to *in-vivo*, in which an afterload threshold must be overcome before tissue shortening can occur.^22^ It is unknown how these different types of loading regulate specific ion channel expression, or how the changes they induce in cardiomyocytes differ from the results seen here. It would be of great benefit to the field to understand how these different types of loading regulate physiology in unique ways.

## Supplementary Materials

Figure S1: Quantification of nuclear alignment and nuclear aspect ratio in wild type µHM

Figure S2: Quantification of action potential and calcium handling parameters for wild type µHM

Figure S3: Effects of saxitoxin, nifedipine, E4031, and BaCl_2_ on wild type µHM

Figure S4: Quantification of RNA expression

Figure S5: Quantification of Ki67 expression in wild type µHM

Figure S6: Quantification of CX43 Protein Expression in wild type µHM

Figure S7: Western blot of PKP2 and GAPDH in wild type and PKP2^-/-^ iPSCM

Figure S8: Quantification of nuclear alignment and nuclear aspect ratio in PKP2^-/-^ µHM

Figure S9: Quantification of action potential and calcium handling parameters for PKP2^-/-^ µHM

Figure S10: Effects of saxitoxin, nifedipine, E4031, and BaCl_2_ on PKP2^-/-^ µHM

Figure S11: Quantification of Cx43 Protein Expression in PKP2^-/-^ µHM

Table S1: Primers Used for SYBR Green Quantitative qRT-PCR Analysis

Table S2: Antibodies Used

## Materials & Methods

### Stem Cell-Derived Cardiomyocyte Production

Wild Type C” human induced pluripotent stem cells (iPSC) were created at the Gladstone Institutes of Cardiovascular Disease (Coriell Institute GM25256), and were modified via knock-in of a single copy of GCaMP6f into the AAVS1 “safe harbor locus”.^26^ iPSC were cultured at 37°C in Essential 8 media on 6-well plates coated with growth factor-reduced Matrigel. Once 85% confluency was reached, the cells were passaged into new wells using Accutase. iPSC were differentiated into cardiomyocytes using small molecule manipulation of Wnt signaling.^36^

To generate PKP2^-/-^ iPSC, the wild-type C iPSC used were modified by CRISPR (indel mutations between R185 and V202) by the Genome Engineering and Stem Cell Center at Washington University. The knockout was confirmed by next-gen sequencing (data not shown) and western blot (Fig 5A, B).

### μHM-Forming Stencil Fabrication

Using Solidworks, the stencil mold was created by first extruding a rectangular base several millimeters thick to prevent warping during the polymer crosslinking process, and eventual print removal from the printer. On top of this base an individual dogbone shape is designed and extruded to a height of 1 mm. Using linear patterning we can easily create groups of dogbones, typically clustered in groups of 3 with several millimeters of spacing between groups. This allows us to easily cut and place groups into wells of a 24 well plate. The individual dogbone shapes have a 1 mm x 1 mm knob/square on each end, with a 400 µm wide shaft of variable length. Importantly, dogbones within a cluster should be spaced at least 2 mm apart from one another to prevent a syncytium of tissue from “bridging” between adjacent micro-muscles (Fig. S3.7). The mold was replicated into PDMS using Hydrogel Assisted STereolithographic Elastomer prototyping (HASTE) as previously described.^35^ Briefly, a negative of the 3D-printed mold is first created by casting off agar. Sylgard 184 (Dow Corning) is mixed according to the manufacturer’s instructions, poured onto the agar negative, degassed, and cured at 37°C for 8+ hours to form a PDMS positive of the 3D-printed mold.

The PDMS replicates of the 3D-printed molds were oxidized using air plasma (Harrick Plasma) for 90 seconds at high power. These were then treated with trichloro(1H,1H,2H,2H-perfluorooctyl)silane via vapor deposition to enable PDMS-off-PDMS molding for final stencil creation. Stencil molds with “dogbone” shaped through-holes were formed by pouring Sylgard 184 over the treated PDMS molds, clamped between glass/acrylic plates and cured at 60°C for 4+ hours. Each dogbone shaped through-hole is 1mm thick, and three different dogbone sizes were created: 1) a tissue that is just a 1 mm x 1 mm knob with no shaft (“0 mm”), 2) a tissue with 1 mm x 1 mm knobs and a 400 µm x 1 mm shaft (“1 mm”), and 3) 1 mm x 1 mm knobs and a 400 µm x 2 mm shaft (“2 mm”) (Fig. 3.1B). Here we utilized stencils of varying shaft length (0, 1, & 2mm long), hypothesized to modulate tissue prestress alongside increasing shaft length without the need to dynamically modulate tissue length.

### Finite Element Modeling

A COMSOL model (Fig. 3.1B) comprised of a linear elastic material confined to the dogbone shape was used, with the material adherent to the substrate only in the knob ends. With an initial strain of –0.5 for each axis (x, y, z), isotropic shrinkage of the µHM volume by 82.5% was performed, compared with initial 100% dogbone-like volume. This modeling predicted stepwise increases in tissue prestress within the center of the tissue shaft. These parameters were chosen based on prior experimental observations on µHM formation and compaction after initial cell seeding. During tissue seeding the single-cell cardiomyocyte/media suspension fills the entirety of the dogbone-shaped stencil. After approximately 24 hours, the cells have self-compacted and assembled to form the µHM. After tissue compaction, the tissue remains adherent to the substrate, in this case tissue culture plastic, while the shaft region is lifted and non-adherent to the substrate nor the PDMS stencil. Based on observations of the µHM before and after tissue formation, it appears the cells compact isotropically.

### μHM Formation

To seed the PDMS stencils, they were cut, dipped in methanol, and placed into the wells of tissue culture well plates. The well plate was then placed into a 60°C oven overnight to reversibly bond the PDMS stencil to the well surface. The well was then coated with fibronectin to enable cardiomyocyte attachment to the substrate.

iPSC-CM at day 15 of differentiation were singularized using 0.25% trypsin and seeded at a density of 7.5*10^7 cells/mL at 3 µL per individual µHM (∼225,000 cells per tissue) without exogenous matrix. Seeded cells were incubated at 37°C for 30-60 minutes before adding media to limit cell loss, using DMEM with 20% FBS, 10 µM Y-27632, 150 µg/mL L-ascorbic acid, 4 µg/mL Vitamin B12, and 3.2 µg/mL penicillin. µHM typically began spontaneous beating within 24-48 hours of seeding; upon observing beating tissues the media was changed to RPMI/B-27, 150 µg/mL L-ascorbic acid, 4 µg/mL Vitamin B12, and 3.2 µg/mL penicillin (collectively called R+ media). µHM were then fed R+ media every 2-3 days until termination.

### Immunohistochemistry

Tissues were fixed using increasing concentrations of paraformaldehyde from 1% to 4%.^20, 23^ PDMS stencils were then removed, and tissues were embedded in 1%. The agar-embedded tissues were then cryoprotected using 15% and 30% sucrose, flash frozen in OCT and cryosectioned at 8-15 um. Sectioning was done parallel to the tissue longitudinal axis. Samples were permeabilized with 0.1% Triton-X-100 for 20 minutes and blocked using 5% BSA in 0.1% Triton-X-100 for 45 minutes at room temperature. Primary antibodies were incubated overnight at 4°C, secondary antibodies were incubated at room temperature for 2 hours, and cell nuclei (ThermoFisher Hoechst 33342) were stained at room temperature for 10 minutes. The samples were then protected against photobleaching with ProLong Gold (Invitrogen P36930), and imaged on an Olympus confocal microscope (Fluoview FV1200, Tokyo, Japan). Specific primary antibodies are listed in Table S1.

### Cell Morphology Quantification

All images were quantified using ImageJ. Nuclear aspect ratio was quantified as the nuclear major axis length divided by the minor axis length, where a value of 1 indicates a perfectly spherical nucleus. Nuclear alignment was quantified as the percentage of nuclei whose major axis aligned along the same vector, where a value of 1 indicates the major axis of all nuclei were perfectly parallel to each other. Nuclei with an aspect ratio > 0.9 were assigned an alignment score of 0 as an alignment for a sphere could not be quantified. Cell length and width were quantified by staining the cell membranes with wheat germ agglutinin (ThermoFisher W849).

### Quantitative PCR

Total RNA was purified from tissues according to the manufacturer’s instructions using RNAqueous Total RNA Isolation Kit (Invitrogen AM1912). Additional steps were conducted to inactivate DNases. cDNA was then synthesized from the resulting RNA using oligo(dT)20 and SuperScript III (Invitrogen 18080051).

Genes were quantified by real-time PCR using SYBR Green primers (Table S2; MilliporeSigma) and was conducted on Applied Biosystems Step One Plus. Data analysis was carried out using log 2-fold change normalized to glyceraldehyde-3-phosphate dehydrogenase (GAPDH) gene expression.

### Western Blot

Tissues were lysed in commercial RIPA buffer (Alfa Aesar J63306), with protease inhibitor (Millipore-Sigma P8340) and Triton-X-100 (Fisher BioReagents BP151) added. Samples were incubated with a reducing agent (Invitrogen NP0009) and buffer (Invitrogen B0007) at room temperature for 20 minutes and loaded into 4-20% polyacrylamide gels (Bio-Rad #4561094) and transferred onto a PVDF membrane (Immobilon-P IPVH00005).

Membranes stained for Nav1.5 were blocked overnight at 4°C with 5% milk powder in tris-buffered saline (TBS), and then stained overnight at 4°C using the Nav1.5 primary antibody at a 1:100 dilution in 5% milk powder in TBS with 0.2% Tween-20 (TBS-T). Membranes stained for PKP2 were blocked overnight at 4°C with 5% milk powder in tris-buffered saline (TBS), and then stained overnight at 4°C using the PKP2 primary antibody at a 1:100 dilution in 5% milk powder in TBS with 0.2% Tween-20 (TBS-T). Secondary antibody staining for these two proteins occurred at room temperature for 1 hour in 5% milk powder/TBS-T. All washing steps occurred at room temperature in 5% milk powder/TBS-T. Membranes stained for Cx43 and GAPDH were blocked for 1 hour at room temperature in 5% bovine serum albumin in TBS. Primary and secondary antibody staining for these two proteins occurred for 1 hour at room temperature, with the antibodies diluted in TBS-T. The SuperSignal West Dura substrate was used for detection (Thermo Scientific 34076) on a Syngene PXi, with ImageJ used to analyze the data.

### Optical analysis of action potential waveforms and calcium transients

These cells were previously genetically modified to express GCaMP6f to image calcium handling dynamics,^68^ and Berst-1 voltage sensitive dye^69^ was used to visualize the action potential. Tissues were imaged on a Nikon Eclipse Tsr2 inverted microscope equipped with a Hamamatsu ORCA Flash 4.0 V3 digital CMOS camera and a Lumencor AURA light engine. A Tokai thermal plate was used to maintain µHM temperature during imaging. Videos were imaged at 200-450 fps. The MATLAB Bio-Formats package was used along with custom MATLAB software^70^ to calculate waveform parameters. To calculate conduction velocity, a modified version of the open-source MATLAB software Rhythm 1.0 was used.^71^

### Pharmacology Study

Saxitoxin (SRM NIST, Product # 8642a) was received in a solution of 80% acidified water (pH 3.5)/20% ethanol. Nifedipine (Sigma-Aldrich, Product #N7634) was initially dissolved in dimethyl sulfoxide. E4031 (Alomone Lab, Cat. #E-500) was initially dissolved in MilliQ water. The maximum percentage of DMSO in the total media for tissues treated with nifedipine was 0.02% at 2 µM of drug, which was considered negligible. BaCl_2_ (Sigma 202738) was initially dissolved in MilliQ water to 100 mM.

All drugs were dissolved into phenol red free RPMI1640/B27 at 10X the desired final concentration, and added to tissues by changing 10% of the media to limit shock due to full media changes. Tissues treated with saxitoxin, nifedipine, or E4031 were allowed to equilibrate for 15 minutes at 37°C before imaging, and were tested at the following final concentrations: 10 nM, 100 nM, 500 nM, and 2 µM. Tissues treated with BaCl_2_ were allowed to equilibrate for 35 minutes at 37°C before imaging, and were tested at a final concentration of 0.5 mM BaCl_2_.

### µHM Field Pacing

Fridericia’s formula was used to beat-rate correct waveform parameters. To ensure Fridericia’s formula was relevant for comparing action potential and calcium handling dynamics across variable beat rates for different tissues, field pacing was used for certain tissues at day 15. 1-3 Hz field pacing was applied using a Myo-pacer EP instrument (IonOptics). Two electrodes were positioned into the wells so that the applied voltage traveled through the individual tissues. 30V, 20 ms bipolar pulse trains were applied. This formula was found to accurately convert the measurements up to 3Hz, which was deemed appropriate for our data as no tissues spontaneously beat above approximately 3Hz (Fig. S3.2B).

### Statistical Analysis

GraphPad Prism 8.3.1 was used for statistical analysis. Ordinary one-way ANOVA was used for comparing multiple data groups before performing post-hoc Holm-Sidak mean comparison test. *P* value < 0.05 was considered statistically significant difference. * = p < 0.05;

** = p < 0.005; *** = p < 0.0005. Non-linear curve fitting and determination of EC_50_ were performed using a 3-parameter fit.

## Acknowledgments

We thank Dr. Michael Vahey for use of his confocal microscope, Dr. Guy Genin for advice on finite element modeling, Dr. Jon Silva for his advice on pharmacologic inhibition of action potential ion channels, Rebecca Mellor and Drs. Stacey Rentschler and Jeanne Nerbonne for advice on Western blot analysis of sodium channels and Cx43 along with positive control lysates from adult mouse ventricular tissues, and members of the Dr. Elizabeth Haswell lab for help with western blot imaging.

## Author contributions

Conceptualization: DWS, DRS, NH

Methodology: DWS, DRS, NH

Investigation: DWS, DRS, JG, KO, GR, MKM, BK, MP, NH

Visualization: DWS, NH Writing—original draft: DWS, NH

Writing—review & editing: DWS, DRS, JG, KO, GR, MKM, BK, MP, NH

Supervision and Funding: NH

## Data and materials availability

All data are available in the main text or the supplementary materials.

## Ethics Approval

Ethics approval not required.

