## Supplemental Data for "Chronic Prestress Regulation of Micro-Heart Muscle Physiology"


This PDF file includes:


Figure S1: Quantification of nuclear alignment and nuclear aspect ratio in wild type μHM

Figure S2: Quantification of action potential and calcium handling parameters for wild type μHM

Figure S3: Effects of saxitoxin, nifedipine, E4031, and $BaCl_2$ on wild type μHM

Figure S4: Quantification of RNA expression

Figure S5: Quantification of Ki67 expression in wild type μHM

Figure S6: Quantification of CX43 Protein Expression in wild type μHM

Figure S7: Western blot of PKP2 and GAPDH in wild type and $PKP2^{-/-}$ iPSCM

Figure S8: Quantification of nuclear alignment and nuclear aspect ratio in $PKP2^{-/-}$ μHM

Figure S9: Quantification of action potential and calcium handling parameters for $PKP2^{-/-}$ μHM

Figure S10: Effects of saxitoxin, nifedipine, E4031, and $BaCl_2$ on $PKP2^{-/-}$ μHM

Figure S11: Quantification of Cx43 Protein Expression in $PKP2^{-/-}$ μHM

Table S1: Antibodies Used

Table S2: Primers Used for SYBR Green Quantitative qRT-PCR Analysis

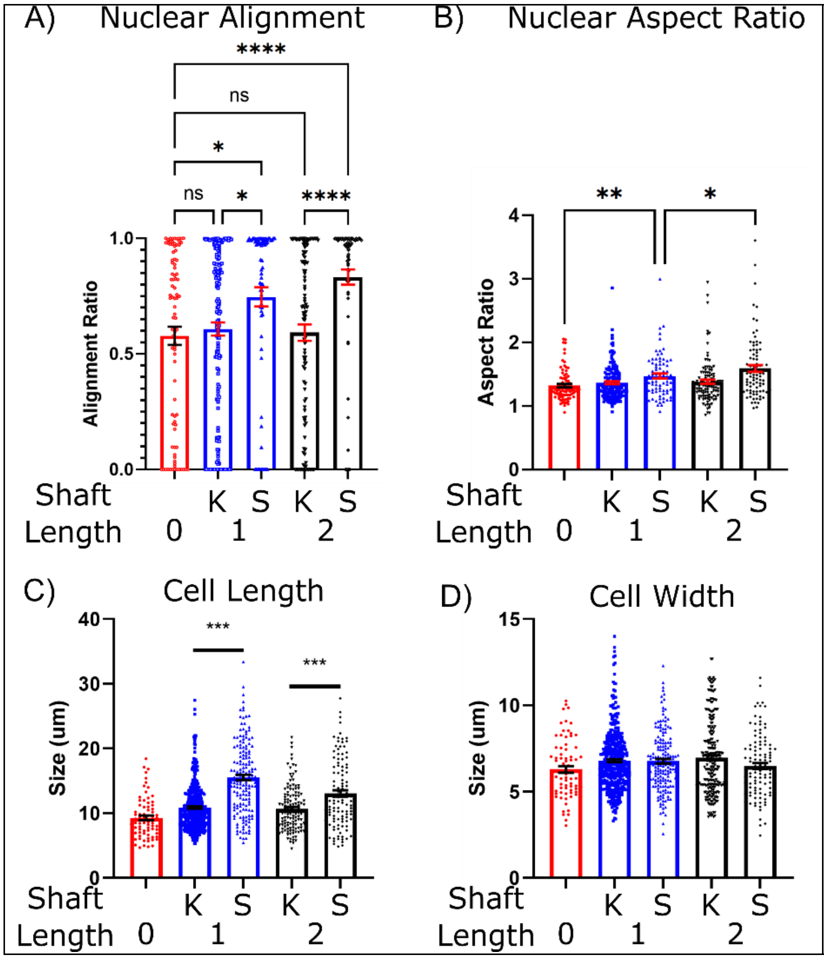


**Supplemental Figure 1**

**A)** Quantification of nuclear alignment and **B)** nuclear aspect ratio at μD15 for 0mm, 1mm, and 2mm μHM. Data for 1mm and 2mm μHM are separated between cells located in the knob (K) and shaft (S) regions of the μHM. **C)** Quantification of cell length and **D)** cell width at μD15 for 0mm, 1mm, and 2mm μHM. Data for 1mm and 2mm μHM are separated between cells located in the knob (K) and shaft (S) regions of the μHM. (*n* > 100 cells from 6-8 μHM, *: $p < 0.05$, **: $p < 0.005$, ***: $p < 0.0005$, error bars: *SEM*)

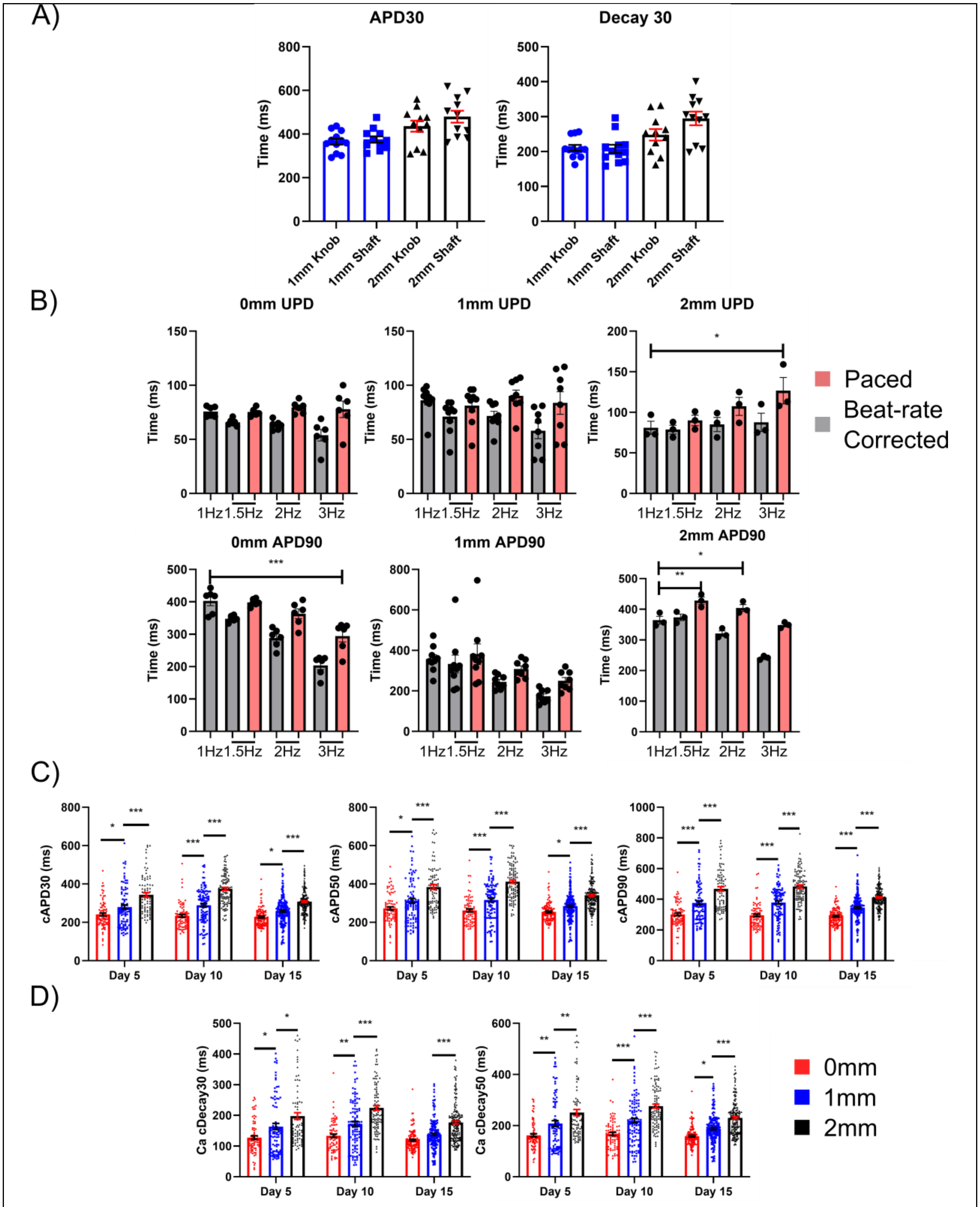


**Supplemental Figure 2 A)** Quantification of action potential amplitude, action potential duration 30, and calcium decay 30 for 1mm and 2mm µHM, comparing these parameters based on where in the tissue they are measured (knob of shaft region). Values were found to be equivalent irrespective of where imaging occurs on the tissue. This justifies the use of an ROI encompassing the entire tissue as used for quantification of electrophysiology as conducted throughout the manuscript. (*n* = 11 μHM) **B)** Quantification of action potential upstroke duration and action potential duration 90 for 0mm, 1mm, and 2mm μHM at multiple pacing frequencies to determine accuracy of Fridericia's formula for beat-rate correction of parameters for these tissues. (*n* = 3-9 μHM) **C)** Quantification of the action potential and **D)** calcium handling parameters that are visualized in the electrophysiology heatmap, for 0mm, 1mm, and 2mm μHM at μD5, μD10, and μD15. (*n* > 100 μHM, *: $p < 0.05$, **: $p < 0.005$, ***: $p < 0.0005$, error bars: *SEM*).

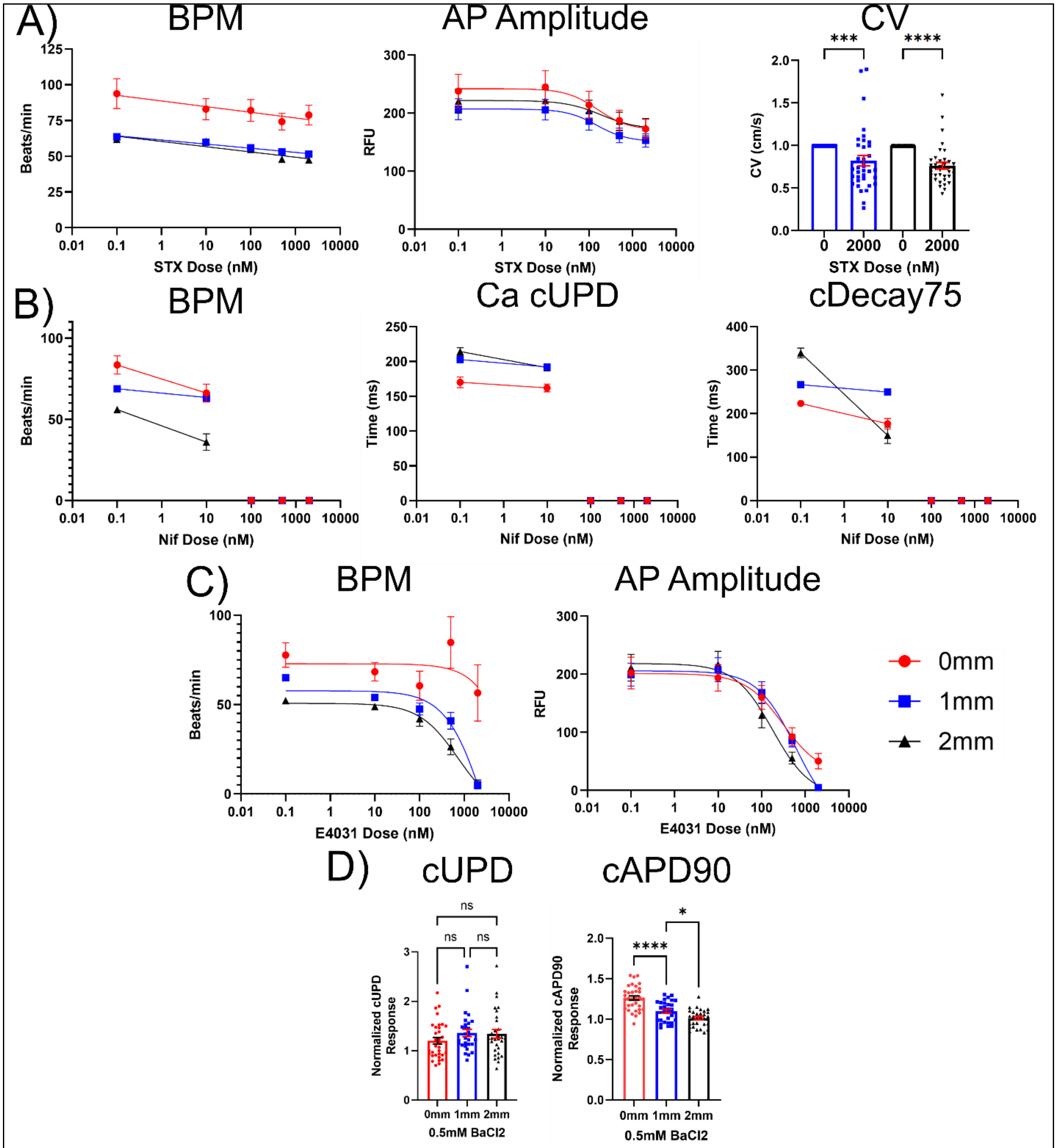


**Supplemental Figure 3 A)** Effects of saxitoxin on µHM spontaneous beat rate, action potential amplitude, and normalized spontaneous conduction velocity at µD15. (*n* = 29-39 µHM) **B)** Effects of nifedipine on µHM spontaneous beat rate, beat-rate corrected calcium transient upstroke duration, and beat-rate corrected decay time 75 at µD15. (*n* = 30-48 µHM) **C)** Effects of E4031 on µHM spontaneous beat rate and action potential amplitude at µD15. (*n* = 26-40 µHM) **D)** Effects of $BaCl_2$ on µHM beat-rate corrected upstroke duration and beat-rate corrected action potential duration 90 at µD15. (*n* = 29-33 µHM) (*: $p < 0.05$, **: $p < 0.005$, ***: $p < 0.0005$, error bars: *SEM*).

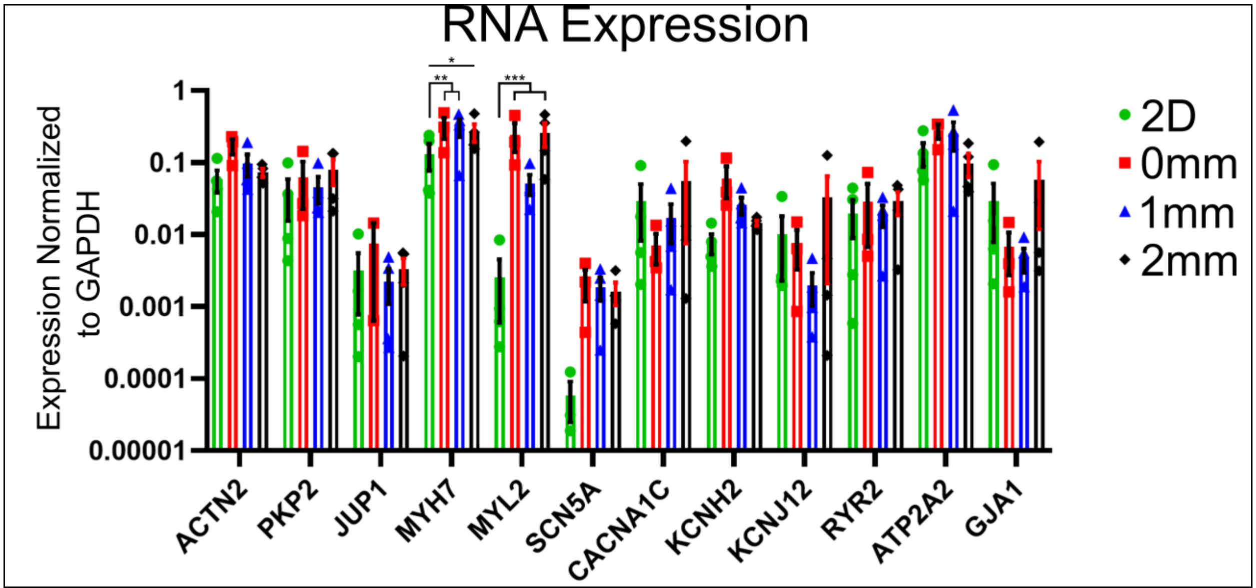


**Supplemental Figure 4** Quantification of RNA expression for 2D (monolayer) cardiomyocytes at D15, and 0mm, 1mm, and 2mm µHM at µD15 for: ACTN2, PKP2, JUP1, MYH7, MYL1, SCN5A, CACNA1C, KCNH2, KCNJ12, RYR2, ATP2A2, and GJA1. (*n* = 4 experimental groups, 6-10 µHM combined per group, *: p < 0.05, **: p < 0.005, ***: p < 0.0005, error bars: *SEM*).

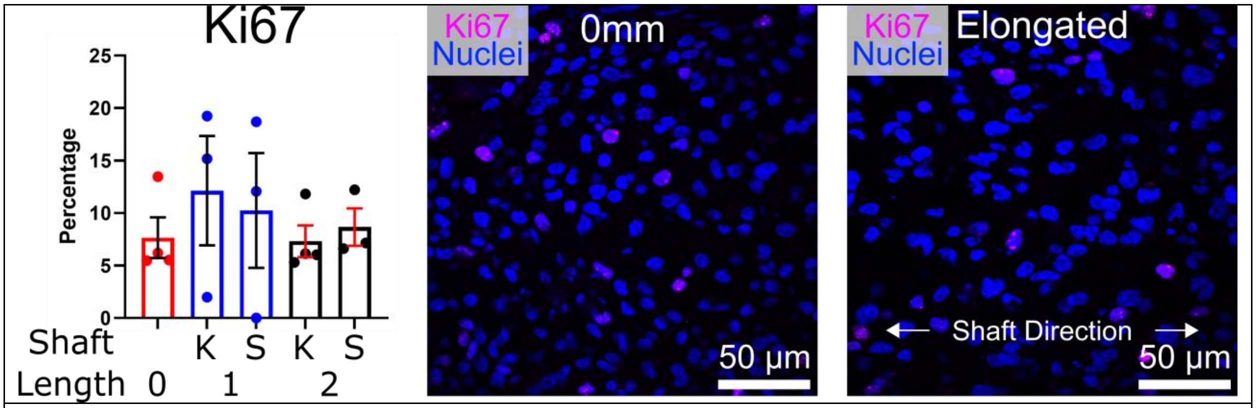


**Supplemental Figure 5 Left)** Quantification of Ki67 expression in 0mm, 1mm, and 2mm µHM at µD15. Data for 1mm and 2mm µHM are separated between cells located in the knob (K) and shaft (S) regions of the µHM. **Right)** Representatives images of Ki67 expression in 0mm and elongated µHM at µD15. (*n* = 3 µHM, error bars: *SEM*).

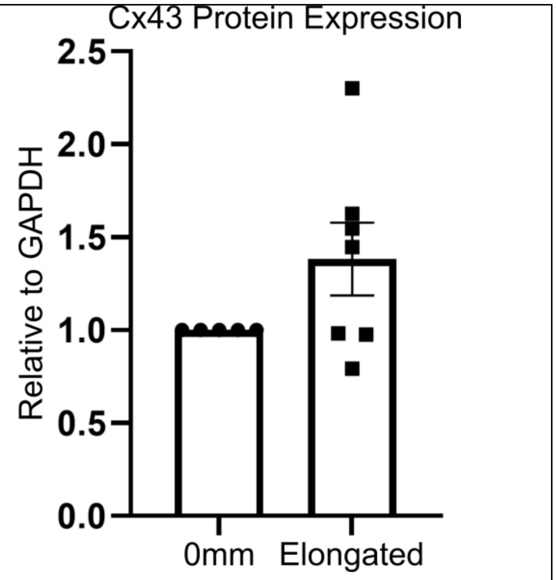


**Supplemental Figure 6** Quantification of CX43 Protein Expression. Each data point represents pooled μHM from a single iPS-CM differentiation batch. Expression is normalized to the value obtained from 0mm μHM from the same biological replicate differentiation ($n \geq 5$).

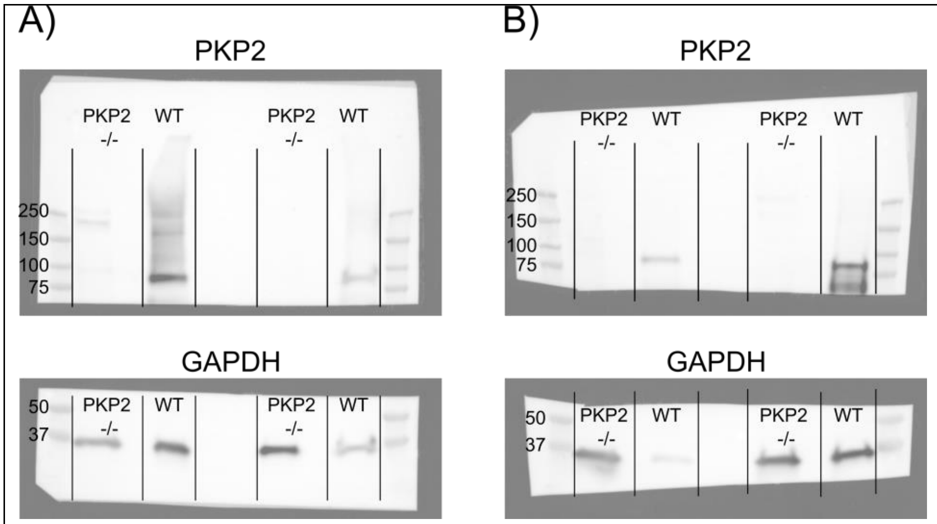


**Supplemental Figure 7 A, B)** Western blot imaging of PKP2 (molecular weight: 92 & 97 kDa) and GAPDH (molecular weight: 36 kDa) in wild type (WT) and $PKP2^{-/-}$ iPSCM, created from 4 different differentiations for both genotypes (2 on each membrane).

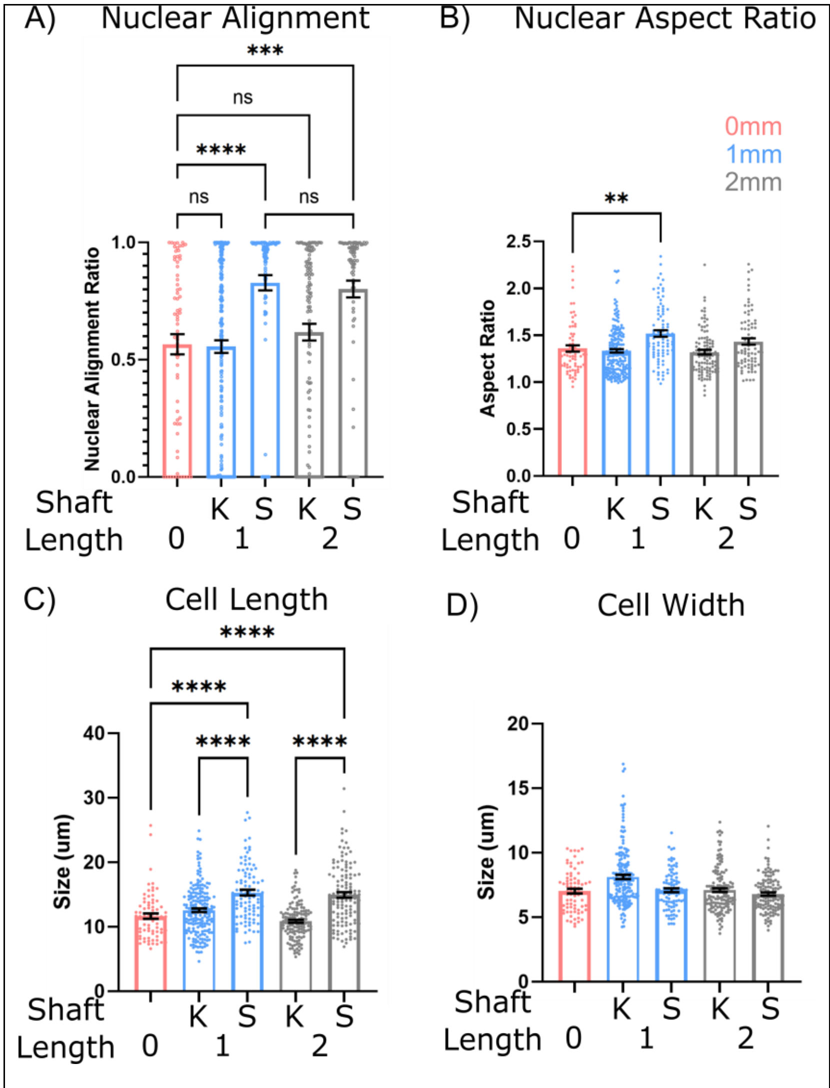


**Supplemental Figure 8**

**A, B)** Quantification of the effects of tissue geometry on nuclear alignment and nuclear aspect ratio in PKP2$^{-/-}$ μHM at μD15. Data for 1mm and 2mm μHM are separated between cells located in the knob (K) and shaft (S) regions of the μHM. (*n* > 70 nuclei from 6-8 μHM per size) **C, D)** Quantification of the effects of tissue geometry on cell length and width in PKP2$^{-/-}$ μHM at μD15. Data for 1mm and 2mm μHM are separated between cells located in the knob (K) and shaft (S) regions of the μHM. (*n* > 100 cells from 6-8 μHM, *: $p < 0.05$, **: $p < 0.005$, ***: $p < 0.0005$, error bars: *SEM*)

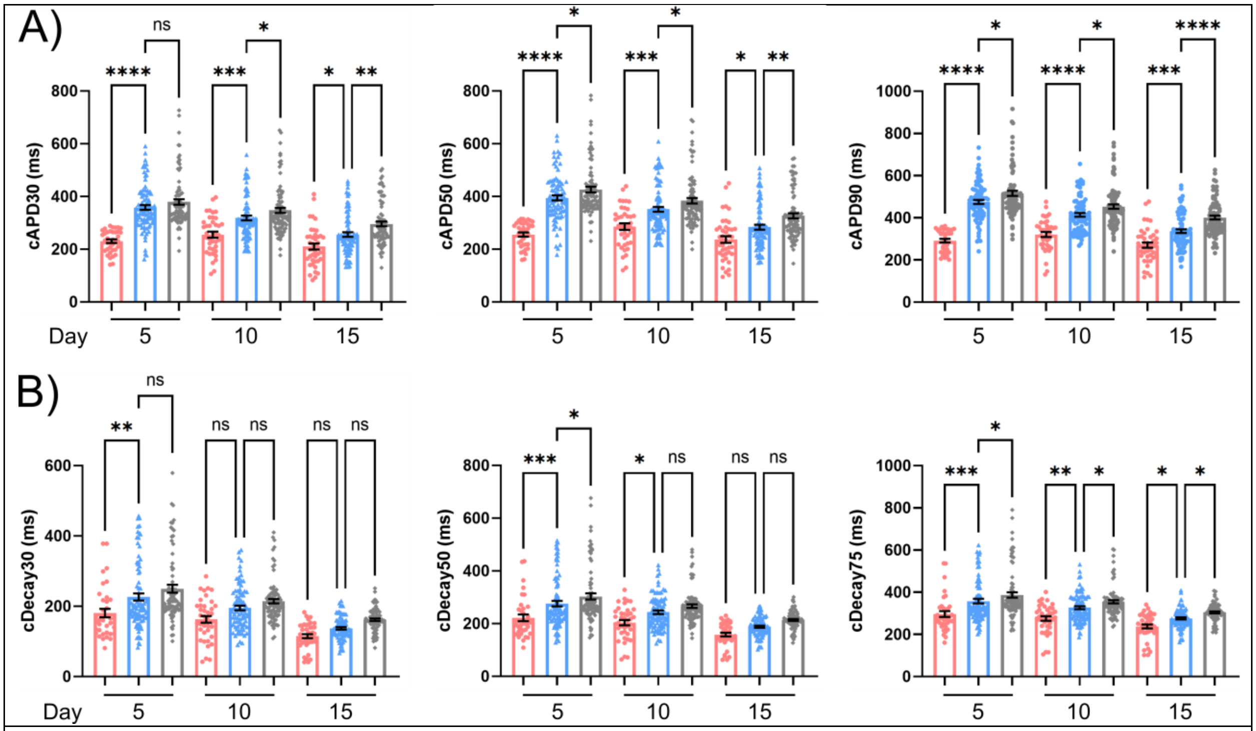


**Supplemental Figure 9 A)** Quantification of additional action potential and **B)** calcium handling parameters for 0mm, 1mm, and 2mm PKP2$^{-/-}$ μHM at μD5, μD10, and μD15. (*n* > 100 μHM, *: p < 0.05, **: p < 0.005, ***: p < 0.0005, error bars: *SEM*).

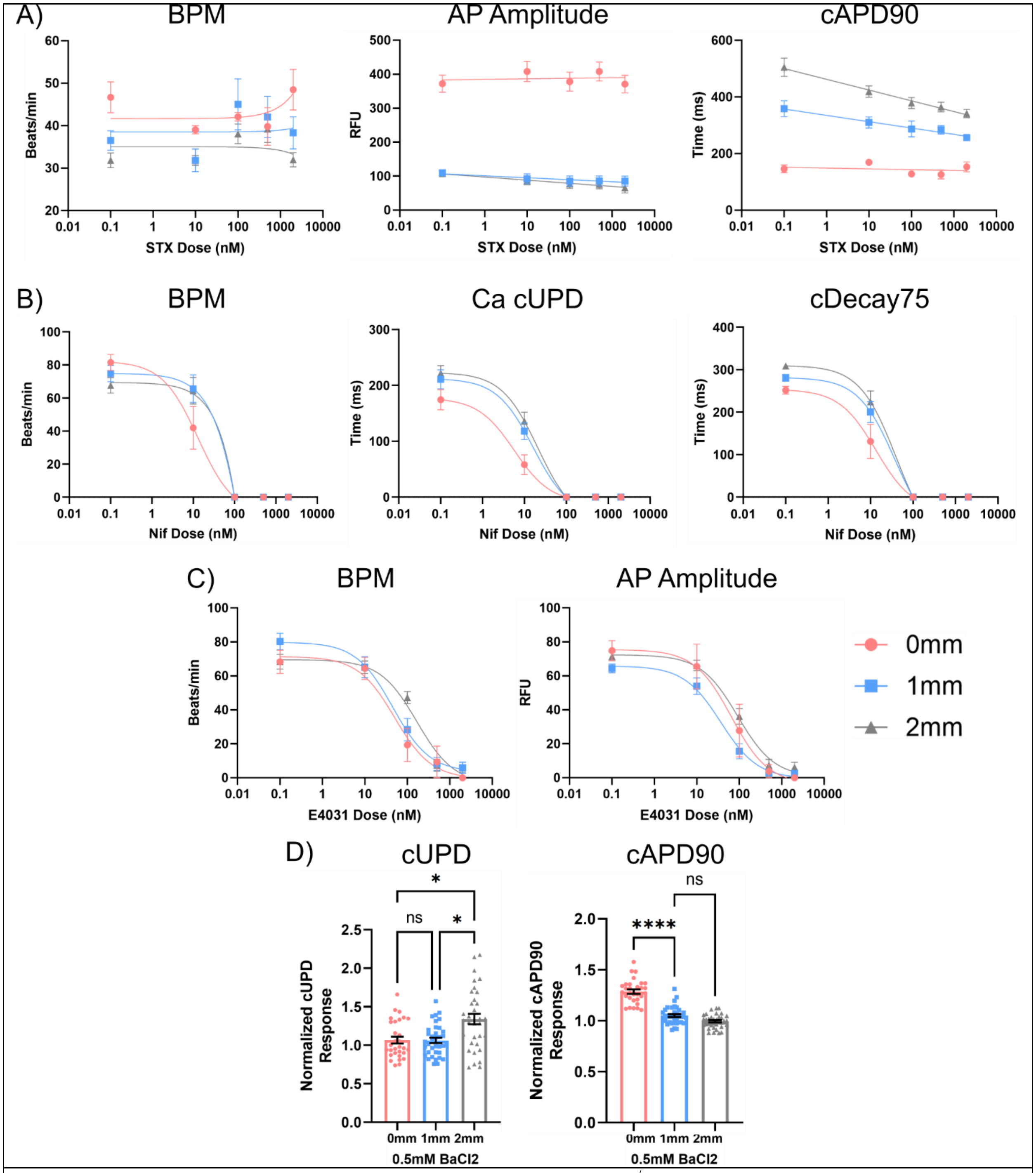


**Supplemental Figure 10** Effects of **A)** saxitoxin on PKP2$^{-/-}$ µHM spontaneous beat rate, action potential amplitude, and beat-rate corrected action potential duration 90 at µD15. (*n* = 3-13 µHM) **B)** nifedipine on PKP2$^{-/-}$ µHM spontaneous beat rate, beat-rate corrected calcium transient upstroke duration, and beat-rate corrected decay time 75 at µD15. (*n* = 12-24 µHM) **C)** E4031 on PKP2$^{-/-}$ µHM spontaneous beat rate and action potential amplitude at µD15. (*n* = 9-23 µHM) and **D)** $BaCl_2$ on PKP2$^{-/-}$ µHM beat-rate corrected upstroke duration and beat-rate corrected action potential duration 90 at µD15. (*n* = 29-33 µHM) (*: p < 0.05, **: p < 0.005, ***: p < 0.0005, error bars: *SEM*).

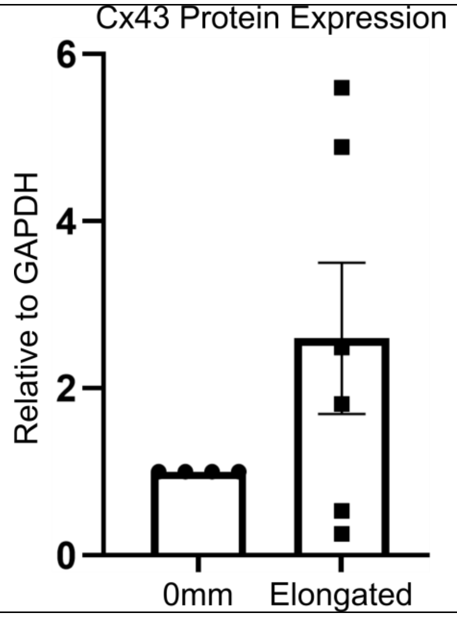


**Supplemental Figure 11** Quantification of Cx43 Protein Expression in PKP2$^{-/-}$ µHM. Each data point represents pooled µHM from a single iPS-CM differentiation batch. Expression is normalized to the value obtained from 0mm µHM from the same biological replicate differentiation ($n \geq 4$).

**Table S1. Antibodies Used**

| Antibody | Product Information | Usage | Dilution and Application | Species |
|---|---|---|---|---|
| Sarcomeric α-actinin (ACTN2) | Sigma EA-53 | IHC | 1:1000 IHC | Mouse |
| Nav1.5 | Cell Signaling D9J7S | IHC | 1:1000 | Rabbit |
| Ki67 | ThermoFisher PA5-19462 | IHC | 1:1000 | Rabbit |
| Cx43 | Sigma-Aldrich C6219 | IHC/WB | 1:1000 | Rabbit |
| PKP2 | Fitzgerald 10R-P130B | IHC | 1:1 | Mouse |
| PKP2 | Abcam ab151402 | WB | 1:100 | Mouse |
| Nav1.5 | Alomone Lab ASC-005 | WB | 1:100 | Rabbit |
| GAPDH | Cell Signaling 14C10 | WB | 1:1000 | Rabbit |
| HRP secondary | Invitrogen 31460 | WB | 1:2500 | Goat anti-Rabbit |
| HRP secondary | Invitrogen 31430 | WB | 1:2500 | Goat anti-Mouse |

**Table S2. Primers Used for SYBR Green Quantitative qRT-PCR Analysis**

| Target gene | 5' → 3' Primer sequences | |
|---|---|---|
| ACTN2 | Forward: CTGCTGCTTTGGTGTCAGAG | Reverse: TTCCTATGGGGTCATCCTTG |
| PKP2 | Forward: ACAAATAACAGGTTTGCTGTGGA | Reverse: TTCAGGCCACCCAGAAAAGG |
| JUP1 | Forward: TCGGTTACTGAGTTGCTGCC | Reverse: CATCGTGGCTACTGCGCC |
| MYH7 | Forward: CGACCTTCTTCTCTTGCTC | Reverse: GAGGACAAGGTCAACACCCT |
| MYL2 | Forward: TACGTTCGGGAAATGCTGAC | Reverse: TTCTCCGTGGGTGATGATG |
| SCN5A | Forward: GAGCTCTGTCACGATTTGAGG | Reverse: GAAGATGAGGCAGACGAGGA |
| CACNA1C | Forward: GGAGAGTTTTCCAAAGAGAG | Reverse: TTTGAGATCCTCTTCTAGCTG |
| KCNH2 | Forward: CAC CGC CCT GTA CTT CATCT | Reverse: AGG CCT TGC ATA CAG GTT CA |
| KCNJ12 | Forward: TGG ATC CTT TCC AGT TGG TG | Reverse: CGG CTC TTG AGT TCT ATCTT |
| RYR2 | Forward: CTGGTGAGGAAGAAGCCAAG | Reverse: TGTCTTCCTGGCTGTGAGTG |
| ATP2A2 | Forward: ACCCACATTCGAGTTGGAAG | Reverse: CCAACGAAGGTCAGATTGGT |
| GJA1 | Forward: CAATCTCTCATGTGCGCTTCT | Reverse: GGCAACCTTGAGTTCTTCCTC |
